# Novel MBLs inhibitors screened from FDA-approved drug library restore the susceptibility of carbapenems to NDM-1-harbouring bacteria

**DOI:** 10.1101/2022.01.16.476523

**Authors:** Yan Guo, Hongtao Liu, Mengge Yang, Rui Ding, Yawen Gao, Xiaodi Niu, Xuming Deng, Jianfeng Wang, Haihua Feng, Jiazhang Qiu

**Affiliations:** Key Laboratory of Zoonosis, Ministry of Education, College of Veterinary Medicine, Jilin University, Changchun, Jilin 130062, China; College of Food Science and Engineering, Jilin University, Changchun, Jilin 130062, China

**Keywords:** Carbapenems, Carbapenem-resistant Gram-negative bacteria, Metallo-β- lactamase, FDA-approved drug library, Drug repurposing

## Abstract

The production of metallo-β-lactamases (MBLs) is one of the major mechanisms adopted by bacterial pathogens to resist carbapenems. One promising strategy to overcome MBLs-mediated carbapenems resistance is to develop effective inhibitors. Repurposing approved drugs to restore the efficacy of carbapenems represents an efficient and cost-effective approach to fight infections caused by carbapenem resistant pathogens. Here, twelve FDA-approved compounds were screened to neutralize the ability of NDM-1. Among these compounds, dexrazoxane, embelin, candesartan cilexetil (CAN) and nordihydroguaiaretic acid (NDGA) were further demonstrated to inhibit all tested MBLs, and showed an *in vitro* synergistic bactericidal effect with meropenem against MBLs-producing bacteria. Mechanistic studies revealed that dexrazoxane, embelin and CAN are metal ion chelating agents, while the inhibition of NDM-1 by NDGA involves its direct binding with the active region of NDM-1. Furthermore, dexrazoxane, embelin and CAN and NDGA dramatically rescued the treatment efficacy of meropenem in three infection models. Our observations indicated that dexrazoxane, embelin, CAN and NDGA are promising carbapenem adjuvants against MBLs-positive carbapenem resistant bacterial pathogens.

## Introduction

The overuse and misuse of antibiotics has led to the rapid development and dissemination of antimicrobial resistance (AMR), which is a serious threat to public health worldwide. Currently, few therapeutic options are available for the treatment of infections caused by multi-drug resistant (MDR) bacteria, especially “ESKAPE” pathogens (*<u>E</u>nterococcus faecium*, *<u>S</u>taphylococcus aureus*, *<u>K</u>lebsiella pneumoniae*, *<u>A</u>cinetobacter baumannii*, *<u>P</u>seudomonas aeruginosa* and *<u>E</u>nterobacter spp* [1]. It was estimated that AMR will result in 10 million patient deaths per year by 2050 unless active and effective actions are mounted [2].

Carbapenems are β-lactam antibiotics that are commonly used as last-resort drugs for the treatment of serious MDR gram-negative bacterial infections [3]. In the past two decades, the clinical consumption of carbapenems has increased dramatically, which is inevitably accompanied by the emergence and prevalence of carbapenem-resistant strains [4]. In particular, carbapenem-resistant *Enterobacteriaceae* (CRE), which has become a global public threat [5]. The production of an inactivating enzyme, carbapenemase, is the major resistance mechanism of *Enterobacteriaceae* to carbapenems [6]. The large numbers of carbapenemases are divided into three major categories according to their amino acid sequences. Classes A and D carbapenemases (e.g. KPC and OXA-48) are serine β-lactamases (SBLs), which utilize an active serine residue to covalently attack the β-lactam ring [7]. Class B carbapenemases represented by NDM, IMP and VIM are metallo-β-lactamases (MBLs), which require zinc ions for the activation of a water nucleophile to hydrolyse the ring [7, 8]. The global spread of MBLs is particularly problematic and has aroused significant concerns due to their ability to inactivate almost all clinically approved β-lactams except for aztreonam [9]. In addition, MBLs have great potential for horizontal gene transfer between various bacterial species through MBLs-bearing plasmids [8]. The SENTRY antimicrobial surveillance program reported that the detection of MBL genes in CRE isolates were rapidly increased from 4.3% during 2007-2009 to 12.7% during 2014-2016, among which NDM was the most predominant type of MBLs, accounting for approximately 10% of the CRE isolates [10].

Carbapenem-resistant bacteria including *Acinetobacter baumannii*, *Pseudomonas aeruginosa* and *Enterobacteriaceae* were listed by the World Health Organization (WHO) in 2019 as the priority pathogens required for new treatment [10, 11]. Compared with the development of novel antibiotics, the discovery of carbapenemase inhibitors represents a fast and cost-effective alternative approach to restore the susceptibility of bacteria to carbapenems [12]. The advantages were further strengthened by advances in rapid detection technologies, enabling fast and accurate detection of carbapenemase genes [13–15]. Indeed, this strategy has yielded the successful development of SBL inhibitors. For example, vaborbactam was approved in 2017 for clinical use in combination with meropenem to overcome carbapenem resistance mediated by the expression of SBLs [16]. However, to date, no MBL inhibitors have been approved for clinical use to date.

Here, we screened 1515 FDA-approved drugs through the nitrocefin hydrolysis assay, resulting in the identification of 12 compounds that are capable of inhibiting NDM-1 activity. We further confirmed that dexrazoxane, embelin, candesartan cilexetil (CAN) and nordihydroguaiaretic acid (NDGA) showed significant synergistic antibacterial effects when used in combination with meropenem against NDM-1 positive bacterial strains. In addition to affecting NDM-1, these compounds also exhibit broad inhibitory effects on other major types of MBLs, including IMP and VIM. Further mechanistic studies revealed that dexrazoxane, embelin and CAN are metal ion chelators, while the inhibition of MBLs by NDGA involves NDGA binding to the active region of NDM-1, preventing the binding of NDM-1 to its substrate and thereby inhibiting the activity of NDM-1. Finally, the combination therapy of the compounds with meropenem restored the treatment efficacy of meropenem in mice infected with NDM-1 harbouring *E. coli* isolates. Taken together, our study provides additional options for the treatment of infections caused by MBLs-positive bacterial pathogens.

## Materials and Methods

### Bacterial strains, plasmids, culture methods and reagents

The bacterial strains, plasmids and primers used in this study are listed in Supplementary **Table S1**, **Table S2** and **Table S3**, respectively. All bacterial strains were grown on Luria-Bertani (LB) plates or in LB broth. All NDM-1-positive strains were originated from our previous studies [17]. *E. cloacae* 20710 and *E. cloacae* 20712 were provided by Dr. Yonghong Xiao at Zhejiang University [18]. *A. baumannii* 21 was provided by Dr. Zhimin Guo at the First Hospital of Jilin University.

The full gene sequences of *bla*_IMP-1_, *bla*_VIM-1_, *bla*_KPC-2_ and *bla*_OXA-48_ were synthesized by GeneScript (Nanjing, China) and were inserted into pET28a or pETSUMO for protein expression.

The FDA-approved drug library was purchased from ApexBio Technology (Cat#L1021, Houston, TX, USA). Dexrazoxane, embelin, CAN and NDGA were dissolved in DMSO (Sigma-Aldrich, St. Louis, MO, USA). Meropenem, amoxicillin, ciprofloxacin, imipenem, erythromycin, chloramphenicol, tetracycline and gentamicin were obtained from Dalian Meilun Biotechnology Co., Ltd. (Dalian, China).

### Protein expression and purification

Plasmids derived from pET28a or pETSUMO for protein production were transformed into *E. coli* strain BL21 (DE3). Transformed *E. coli* was inoculated into LB broth containing kanamycin (final concentration of 30 μg/mL) and grown to an OD_600nm_ of 0.8 at 37 °C, followed by induction with isopropyl-β-d-thiogalactoside (IPTG) overnight at 18 °C. Cells were collected by centrifugation and lysed by a homogenizer (JN-mini, JNBIO, Guangzhou, China). Lysed samples were centrifuged at 12, 000 rpm for 20 min. NDM-1 and its mutant proteins, IMP-1, VIM-1, KPC-2 and OXA-48 were purified by nickel affinity chromatography. After washing with lysis buffer, proteins were eluted with 250 mM imidazole and dialyzed twice in a buffer containing 20 mM Tris-HCl (pH 7.5), 150 mM NaCl, 10% glycerol and 1 mM dithiothreitol (DTT). Protein concentrations were measured by the Bradford Protein Assay (Bio-Rad).

### Nitrocefin hydrolysis assays

Nitrocefin hydrolysis assays were determined as described previously [17] in the presence of the indicated concentrations (from 0 to 64 μg/mL) of different compounds. The results were read in 96-well plates at 37 °C using a microplate reader at 492 nm (SYNERGY H1, BioTek). The inhibitory effects of the identified active compound on other carbapenemases (NDM-3, NDM-9, IMP-1, VIM-1, KPC-2 and OXA-48) were tested as described above using nitrocefin as the substrate.

### Determination of the MIC and FICI

MIC values were determined according to the broth microdilution guidelines of the Clinical and Laboratory Standards Institute (CLSI). After the tested strains were diluted with LB to a final concentration of 5×10^5^ CFUs/mL, various concentrations of meropenem (0-256 μg/mL) and increasing concentrations of inhibitors (0-64 μg/mL) were added to the sterile 96-well plate, and further incubated for 18-24 h at 37 °C. The lowest concentration with no visible growth was considered as the MIC value. The synergistic effect of antibiotics and inhibitors was assessed by determining the fractional inhibitory concentration (FIC) index values according to the formula: FICI = (MIC of inhibitors in combination/MIC of inhibitors) + (MIC of antibiotics in combination/MIC of antibiotics).

### Combined disc tests

Overnight bacterial culture was diluted with LB broth to an OD_600nm_ of 0.1. 200 μL of the bacterial suspension was plated on the LB plates. 10 μL of each inhibitor was added to the discs that containing 10 μg meropenem (Oxoid Ltd. Basingstoke, United Kingdom). The discs were placed in the center of the LB plates and incubated at 37 °C for 24 h. Then, the inhibition zone of different treatments was measured and recorded.

### Time-dependent killing

To determine the *in vitro* time-dependent killing of NDM-1 producing bacteria by meropenem, bacterial strains were incubated with inhibitors (32 or 64 μg/mL), meropenem (2 or 8 μg/mL), or inhibitors (32 or 64 μg/mL) in combination with meropenem (2 or 8 μg/mL) at 37 °C. At the indicated time points, samples of different treatments were collected, diluted and plated on LB plates. The CFU values were calculated after incubation overnight at 37 °C.

### Inductively coupled plasma-mass spectrometry (ICP-MS)

ICP-MS was performed as described earlier [19] to investigate the potential ability of the identified NDM-1 inhibitors to chelate zinc ions. Briefly, in order to remove the contained metals, freshly purified NDM-1 protein was exchanged in ICP-MS buffer (20 mM HEPES, 100 mM NaCl, pH 7.5) overnight at 4 °C in a 15 kDa cutoff dialysis tubing. The NDM-1 concentration was adjusted to 5 mg/mL prior to mixing with different concentrations of inhibitors and incubating for 3 h at room temperature with shaking. The NDM-1-inhibitor samples were then dialyzed overnight at 4 °C with ICP-MS buffer by the 12-14 kDa cutoff D-tube dialyzer mini (EMD Biosciences) microdialysis cassettes. The samples were diluted to 1 mg/mL with ICP-MS buffer and then diluted 40-fold with an internal standard containing 10 ng/mL Sc^45^ and 1% nitric acid. Then, the samples were analysed by ICP-MS (XSERIES 2, Thermo Fisher Scientific).

### Metal ions restoration assays

NDM-1 was pretreated with various concentrations of the inhibitors at 37 °C for 10 min, followed by the addition of metal salts (ZnSO_4_, MgSO_4_ and CaCl_2_) and nitrocefin to a final volume of 200 μL and an incubation for 30 min at 37 °C. Then, the absorbance at 492 nm of each sample was measured to calculate the percent residual activity.

### Molecular dynamics simulation

Molecular dynamics simulations of the NDGA-NDM-1 complexes were performed using the Gromacs 4.5.2 software package [20] based on the GRO MOS96 54a7 force field and TIP3P water model. The molecular mechanics/Poisson-Boltzmann surface area (MM-PBSA) method was used to calculate the binding free energy after simulation as described previously [17].

### Secondary structure determination of NDM-1 by circular dichroism (CD) spectroscopy

The CD spectrum of NDM-1 was analysed using a CD spectrophotometer (MOS- 500; Bio-Logic, France) [21]. 200 μg of NDM-1 was incubated with 32 μg/mL of each inhibitor. Then, the secondary structure of NDM-1 (0.2 mg/mL) was determined at room temperature using a quartz cuvette with an optical distance of 1 mm. The scan wavelength range was 190 to 250 nm with a resolution of 0.2 nm and a bandwidth of 1 nm. The BeStSel web server was used to analyse the secondary structure measurements of each sample.

### Plasmid stability

NDM-1-positive *E. coli*, *K. pneumoniae* and *A. baumannii* strains were grown in LB broth overnight with constant shaking in the presence or absence of inhibitors (32 μg/mL). Bacteria were diluted and plated on blank LB plates or LB plates containing 30 μg/mL of kanamycin and incubated at 37 °C for CFU determination.

### Antibodies and Immunoblotting

Polyclonal antibodies against NDM-1 were produced by immunization of mice with recombinant His_6_-NDM-1 (AbMax Biotechnology Co., Ltd., Beijing, China). NDM-1-positive bacterial strains were cultured in the presence of increasing concentrations of inhibitors at 37 °C for 12 h. Bacterial cultures were centrifuged at 12,000 rpm for 5 min, resuspended in 1× loading buffer and subjected to SDS-PAGE. The proteins were transferred to polyvinylidene fluoride (PVDF) membranes followed by a blocking step using 5% nonfat milk. Membranes were incubated with primary antibody (1: 2000) for 2 h and secondary antibody for 1 h at room temperature. The expression of NDM-1 was detected by the Odyssey® CLx Imaging System (Li-Cor).

### Ethics statement

All animal studies were conducted according to the experimental practices and standards approved by the Institutional Animal Care and Use Committee of Jilin University (ALKT202102001). The laboratory animal usage license number is SYXK-2021-0001, as certified by the Department of Science & Technology of Jilin Province. All surgery was performed under sodium pentobarbital anesthesia, and every effort was made to minimize suffering.

### Mouse infection models

Female BALB/c mice (6-8 weeks old, approximately 20 g) were used in this study. Mice were injected intraperitoneally with *E. coli* ZJ487 at a dose of 2×10^8^ CFUs for survival experiments, intramuscularly injected at a dose of 2×10^7^ CFUs for bacterial burden experiments on thigh muscles or inoculated in the left nare (5×10^7^ CFUs) to generate a pneumonia model. For all experiments, after the bacterial challenge, mice were given subcutaneous injections of DMSO, meropenem (10 mg/kg), inhibitors (80 mg/kg), or a combination of meropenem (10 mg/kg) with inhibitors (80 mg/kg) every 12 h. The infected mice were monitored until 96 h post infection for survival analysis. For the thigh muscle and pneumonia models, mice were euthanized 72 h post infection and the thigh muscles and lungs were harvested. Organs were placed into 1 mL of sterilized PBS and homogenized. Then, the suspension was diluted with PBS and plated on an LB plate for CFU enumeration. In addition, lungs were placed into 4% formalin, stained with haematoxylin and eosin and scanned with a digital slide scanner (Pannoramic MIDI, 3DHISTECH Ltd).

### Statistical analysis

The statistical analysis was performed by GraphPad Prism 5 and SPSS software. All the data are presented as the mean ± SD. For *in vitro* studies, the statistical analysis was calculated by the unpaired two tailed Student’s *t*-tests; for *in vivo* experiments, the statistical significance was determined using the log-rank (Mantel-Cox) test (survival rates) and Mann-Whitney U test (tissue bacterial load). *P* <0.05 was considered as significant difference, with * indicating *P* <0.05 and ** indicating *P* <0.01.

## Results

### Four new NDM-1 inhibitors restore the sensitivity of NDM-1-positive bacterial strains to meropenem

To probe for the potential application of an existing drug as an NDM-1 inhibitor, we screened a commercially available FDA-approved drug library comprising 1515 compounds for their ability to inhibit the hydrolysis of nitrocefin to chromogenic cephalosporin by purified NDM-1 enzyme. The initial screening resulted in the identification of 12 compounds that were capable of inhibiting the enzymatic activity of NDM-1, with IC_50_ values ranging from 1.77 μg/mL to 10.71 μg/mL (**Figure 1** and **Table S4**). None of the compounds showed antibacterial activity against the tested gram-negative bacterial strains, as evident by all the MICs being no lower than 128 μg/mL. We further determined the potential synergistic effects of the compounds with carbapenems. Only four compounds (dexrazoxane, embelin, CAN and NDGA) significantly restored the susceptibility of meropenem against engineered *E. coli* strains expressing NDM-1 (BL21-pET28a-SP-NDM-1) or carbapenem-resistant bacterial isolates harbouring *bla_NDM-1_*, as indicated by an FICI less than 0.5 (**Figure 2A** and **Tables S5-S8**). In addition, the combined disk tests also showed significant synergy between meropenem and the inhibitors. Compared to the disks containing meropenem alone, the diameter of the inhibition zone was significantly larger in disks containing both meropenem and each tested inhibitor (**Figure 2B** and **Figure2-figure supplement 1**). Moreover, the time-dependent killing curves were determined to investigate their synergistic bactericidal activity. We found that the combinations led to almost complete bacterial killing within 4 to 10 h for engineered *E. coli* BL21 expressing NDM-1 as well as the clinical *E. coli* isolate ZJ487 (**Figure 2C**). Thus, our results identified that dexrazoxane, embelin, CAN and NDGA represent effective NDM-1 inhibitors displaying synergistic antibacterial activity against NDM-1-positive bacteria with meropenem.

**Figure 1.**
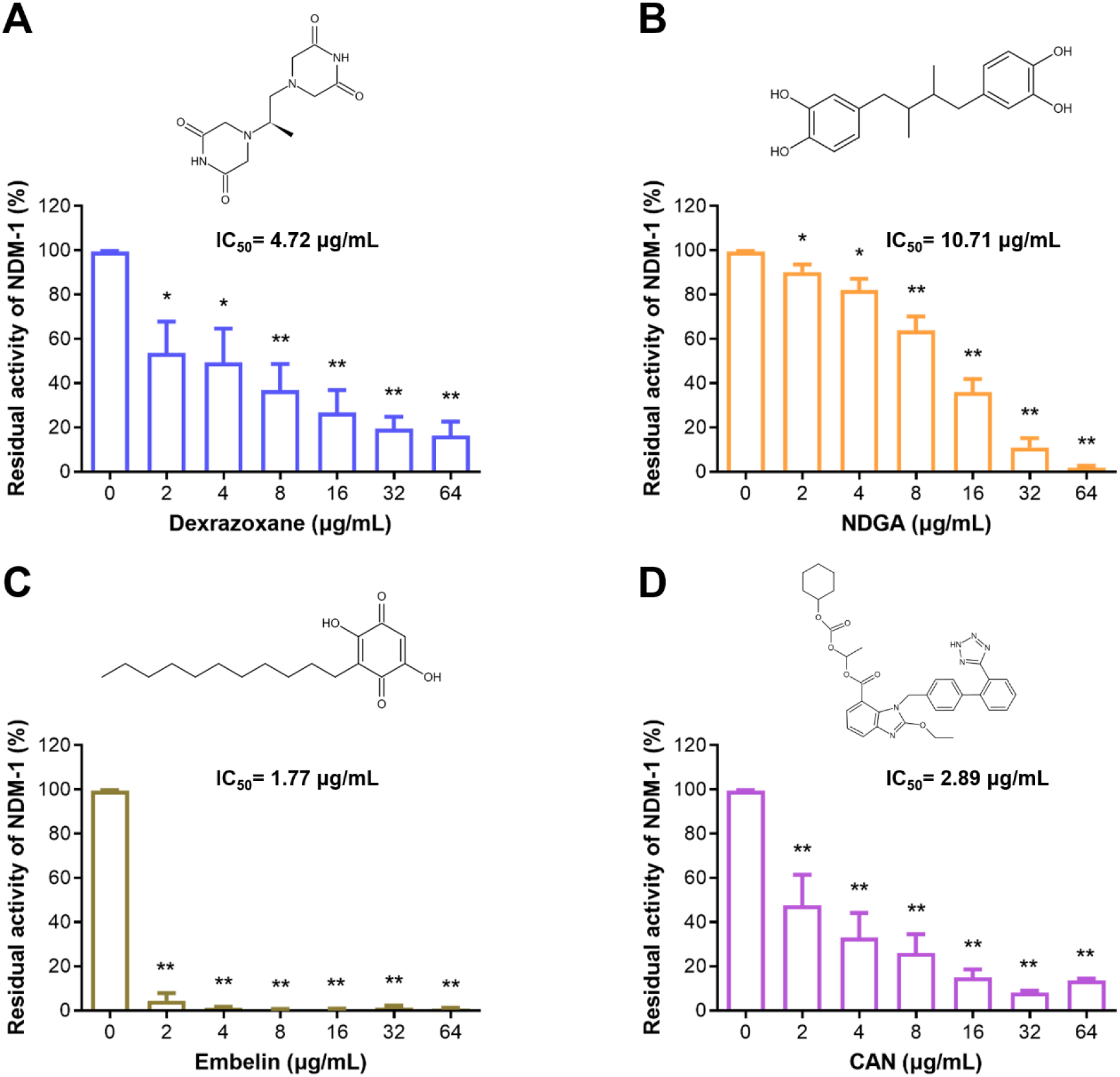
Four inhibitors were screened for their ability to inactivate NDM-1. Inhibition of NDM-1 by dexrazoxane (**A**), NDGA (**B**), embelin (**C**) and CAN (**D**). The chemical structures of the inhibitors are shown in each panel. The positive control was performed in the presence of NDM-1 without inhibitors, while the negative control was carried out in the absence of enzyme. Percent residual activity of NDM-1 = A−A0/A100−A0 × 100%, where A represents the absorbance of samples at 492 nm, A0 and A100 represent 0% and 100% activity of enzyme as determined in the negative control and positive control, respectively. All the data represent the mean ± SD from three independent experiments. * indicates *P* < 0.05 and ** indicates *P* < 0.01 by Student’s *t*-test.

**Figure 2.**
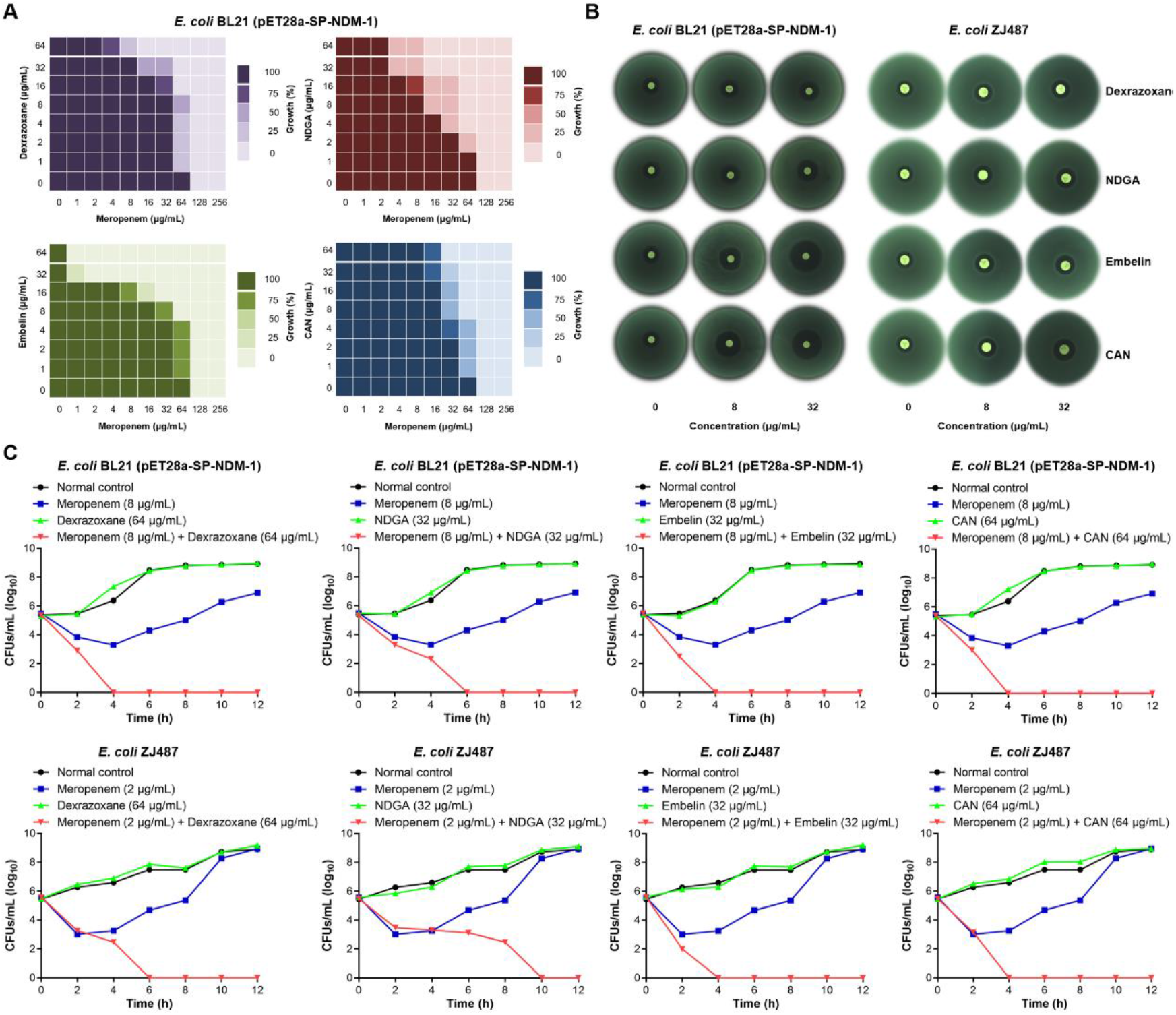
Dexrazoxane, NDGA, embelin and CAN rescue the antibacterial activity of meropenem *in vitro*. (**A**) Microdilution checkerboard analysis showed the synergistic antibacterial effect of dexrazoxane, NDGA, embelin and CAN and meropenem against *E. coli* BL21 (pET28a-SP-NDM-1). (**B**) Zones of inhibition surrounding meropenem disks supplemented with increasing concentrations of dexrazoxane, NDGA, embelin and CAN for the NDM-1-positive strains. (**C**) Time-dependent killing by the combination of meropenem and dexrazoxane, NDGA, embelin or CAN against ZJ487 and *E. coli* BL21 (pET28a-SP-NDM-1). The data shown in panels **A, B** and **C** are one representative of three independent experiments.

### Dexrazoxane, embelin, CAN and NDGA are broad-spectrum MBLs inhibitors

To verify the specificity of the NDM-1 inhibitors identified from the FDA-approved drug library, we purified a series of carbapenemases and tested their enzymatic activity in the presence of dexrazoxane, embelin, CAN and NDGA. Not surprisingly, all the inhibitors displayed potent inhibitory effects on the NDM-3 and NDM-9, the NDM-1 variants that have only a single amino acid mutation (**Figure 3A-B**). The inhibitors also showed increased antibacterial activity of meropenem against clinical isolates expressing NDM-9 (**Tables S5-S8**). Furthermore, the ability of two other major types of MBLs, IMP-1 and VIM-1, to hydrolyse nitrocefin was significantly suppressed by these compounds (**Figure 3C-D**). Therefore, dexrazoxane, embelin, CAN and NDGA could act as broad-spectrum MBLs inhibitors. Indeed, they were able to rescue the susceptibility of meropenem against carbapenem-resistant *E. cloacae* isolates mediated by the production of VIM-1 (**Tables S5-S8**). Surprisingly, embelin, CAN and NDGA but not dexrazoxane also exhibited varying degrees of inhibition of class A (KPC-2) and class D (OXA-48) carbapenemases (**Figure 3- figure supplement 1A-B**). Taken together, these results indicated that dexrazoxane, embelin, CAN and NDGA are broad-spectrum MBLs inhibitors.

**Figure 3.**
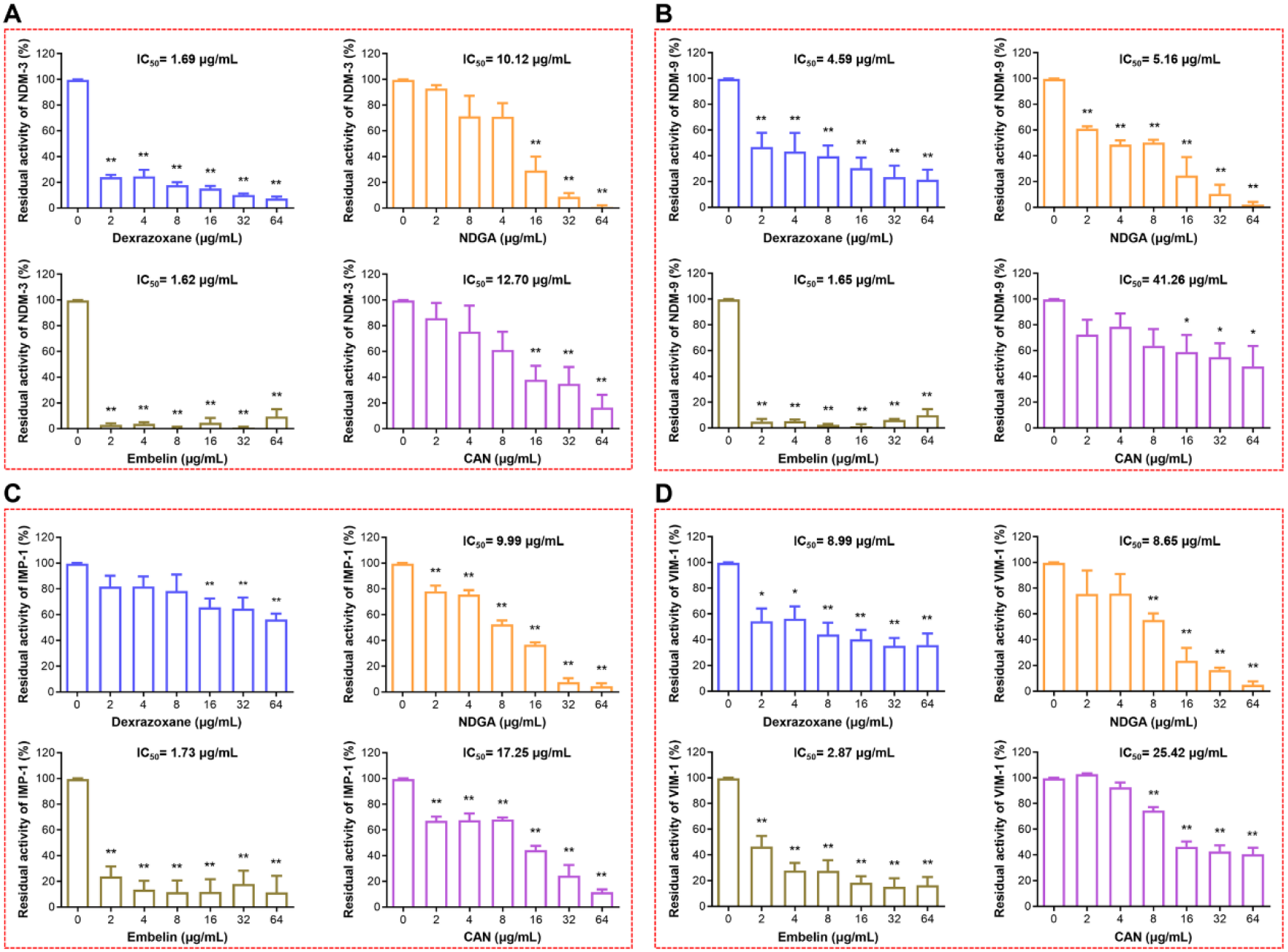
Dexrazoxane, NDGA, embelin and CAN suppress the activity of NDM-3, NDM-9, IMP-1 and VIM-1. Following pre-incubation of dexrazoxane, NDGA, embelin and CAN with NDM-3 (**A**), NDM-9 (**B**), IMP-1 (**C**) and VIM-1 (**D**), the residual enzymatic activity of these proteins was determined as shown in Figure 1. The data shown are the mean ± SD from three independent experiments. * indicates *P* < 0.05 and ** indicates *P* < 0.01 by Student’s *t*-test.

### Dexrazoxane, embelin and CAN act as metal ion chelators to inhibit the activity of NDM-1

To gain insights into the mechanisms adopted by the inhibitors to neutralize NDM-1, we first added excessive zinc ions in the nitrocefin hydrolysis reactions in the presence or absence of inhibitors. The efficacy of dexrazoxane, embelin and CAN in suppressing NDM-1 enzymatic activity was significantly reduced with the addition of 500 μM of ZnSO_4_ (**Figure 4A-B**). Additionally, supplementation with other divalent metal ions (magnesium and calcium ions) in the reaction resulted in a similar restoration of NDM-1 activity (**Figure 4C-D**). Therefore, the inhibition of NDM-1 by dexrazoxane, embelin and CAN may be involved in a metal depletion mechanism. Indeed, after incubation of NDM-1 with 8 or 32 μg/mL of dexrazoxane, embelin and CAN, the zinc ion concentrations associated with NDM-1 were significantly reduced as determined by ICP-MS (**Figure 4E**). Taken together, dexrazoxane, embelin and CAN function as metal chelating agents to suppress the enzymatic activity of MBLs. In contrast, we did not observe either restoration of hydrolysing activity by supplementation with metal ions or the loss of zinc ions in NDM-1 inactivated by NDGA (**Figure 4A-E**), suggesting a different inhibitory mechanism employed by NDGA.

**Figure 4.**
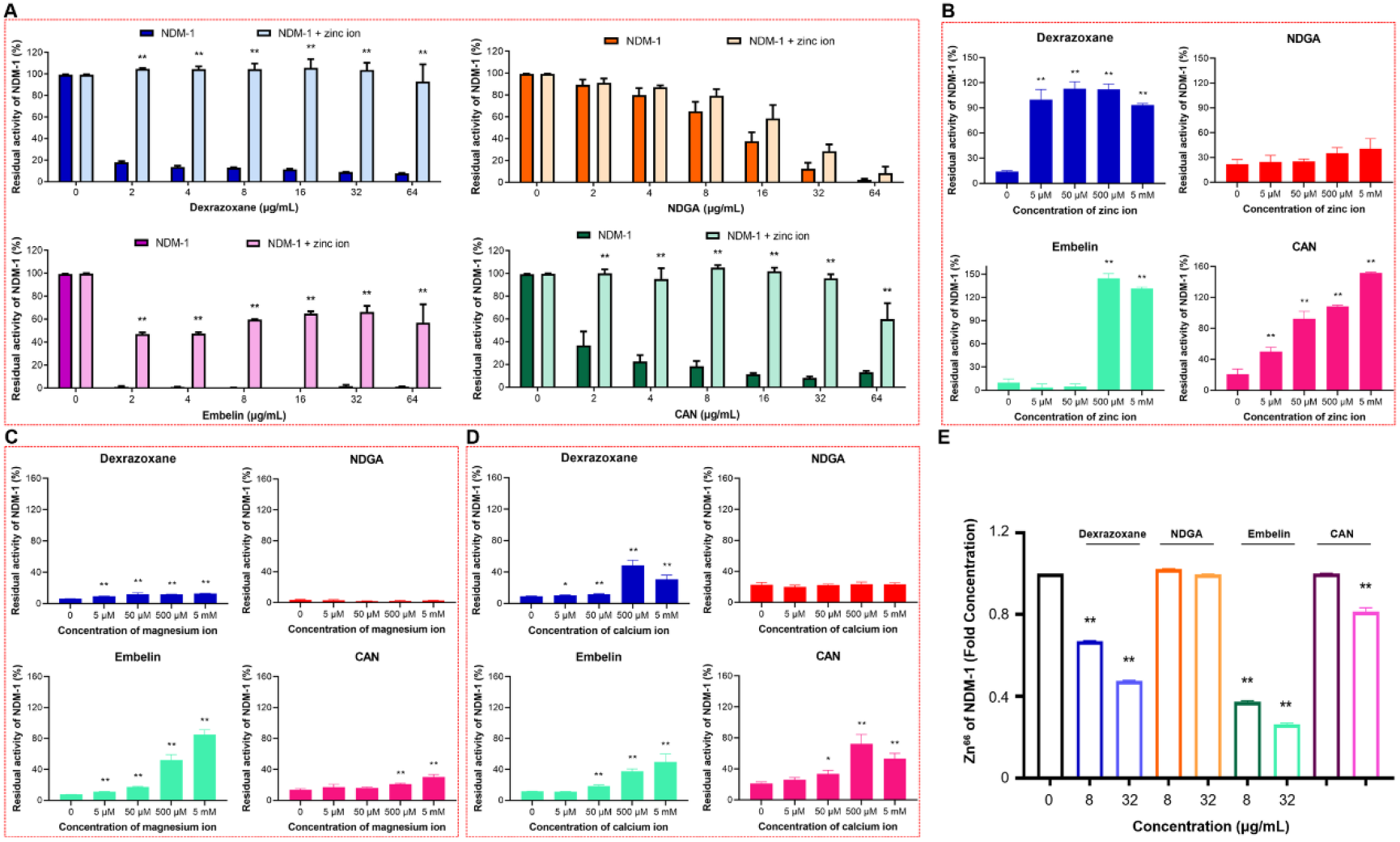
Dexrazoxane, embelin and CAN act as metal ion chelators to inhibit NDM-1 activity. (**A**) Residual activity of NDM-1 after the addition of 500 μM zinc ions in the presence of increasing concentrations of dexrazoxane, NDGA, embelin and CAN. **(B-D)** Supplementation with increasing concentrations of zinc ions **(B)**, magnesium ions **(C)** and calcium ions **(D)** relieved the inhibition of NDM-1 mediated by dexrazoxane, embelin and CAN. (**E**) Zinc ion depletion by dexrazoxane, embelin and CAN as determined by ICP-MS. The vertical coordinate represents the fold concentration of free Zn^66^ in NDM-1. The data shown are the mean ± SD from three independent experiments. * indicates *P* < 0.05 and ** indicates *P* < 0.01 by Student’s *t*-test.

### A direct engagement of NDGA inhibits NDM-1 activity

To further clarify the mechanism of NDGA on the inhibition of NDM-1, we carried out molecular dynamics simulations. A complex MD simulation of the NDM-1-NDGA complex was aimed to explore the binding mode (**Figure 5A**). The root mean square deviation (RMSD) of the complex fluctuated between 0.3 and 0.35 nm after 20 ns, which indicated that the last 80 ns period of the simulation was suitable for the subsequent analysis (**Figure 5B**). Energy decomposition analysis confirmed that the side chains of IIe35, Cys208, Lys211, Asp212, Ala215, Met248 and His250 could bind to NDGA via van der Waals interactions (**Figure 5C**). Moreover, the distance between different residues of NDM-1 and NDGA was analysed. The IIe35, Lys211 and His250 in the NDM-1-binding region are closer to NDGA than other residues (distance < 0.4 nm) (**Figure 5D**). The simulated residues (IIe35, Lys211 and His250) that had the highest binding energy were selected for site-directed mutagenesis. Single mutations of IIe35, Lys211 and His250 into alanine did not affect the enzymatic activity of NDM-1. However, the inhibitory effect of NDGA on NDM-1_I35A_ and NDM-1_K211A_ but not NDM-1_H250A_ was significantly reduced (**Figure 5E**). Collectively, we speculated that NDGA could bind IIe35 and Lys211 to inhibit the activity of NDM-1.

**Figure 5.**
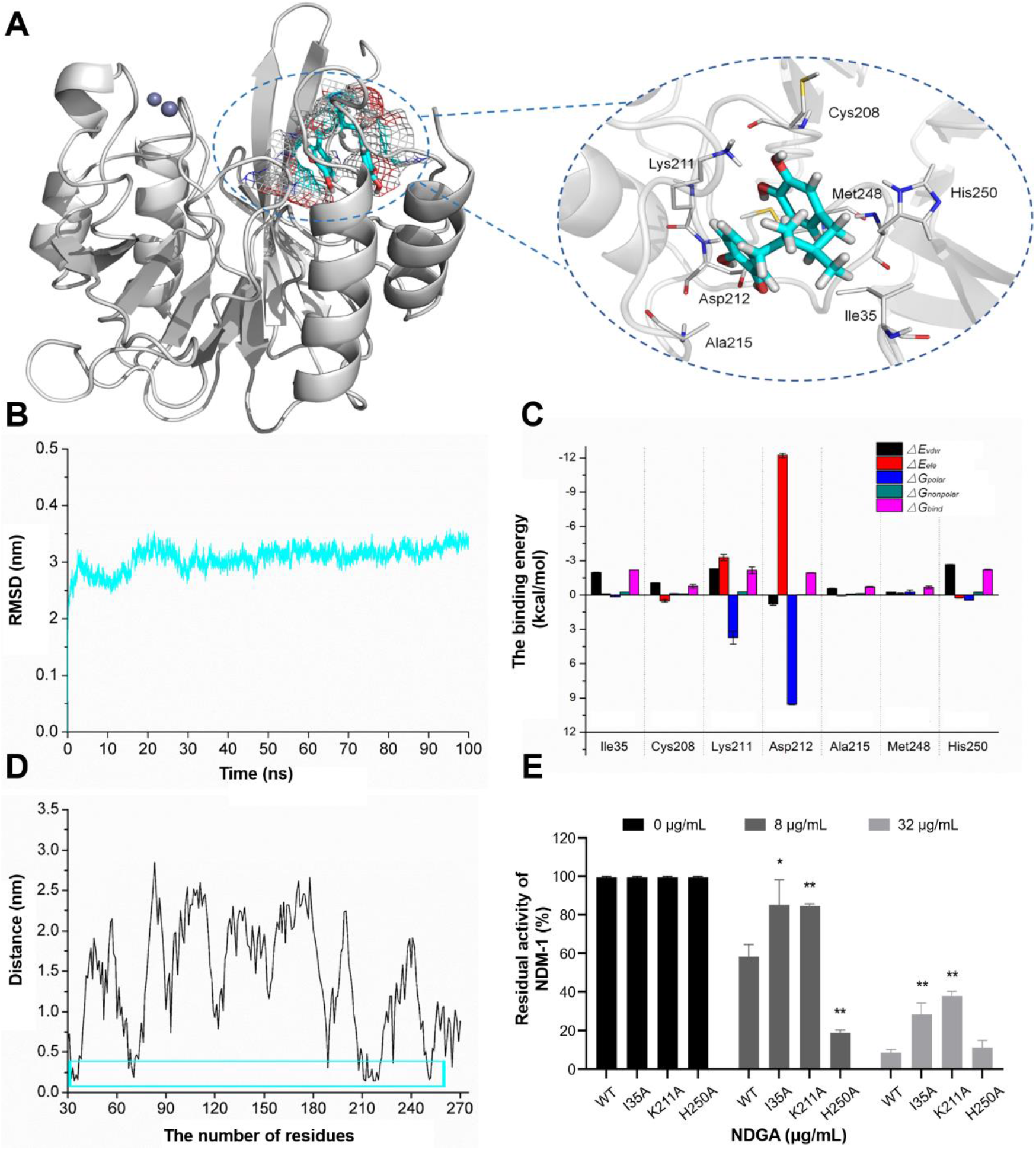
Direct engagement of NDGA with NDM-1. (**A**) The three-dimensional structure determination of NDM-1 with the NDGA complex via a molecular modelling method. The purple spheres represent zinc ions. (**B**) The RMSD values of the NDM-1-NDGA complex. (**C**) Decomposition of the binding energy on a per-residue basis in the binding sites of the NDM-1-NDGA complex. (**D**) Analysis of the distance between all residues of NDM-1 and NDGA. (**E**) Residual activity of NDM-1 and its mutants in the presence of different concentrations of NDGA. Data shown in panel E are the mean ± SD. * indicates *P* < 0.05 and ** indicates *P* < 0.01 by Student’s *t*-test.

### Alteration of the secondary structure of NDM-1 by dexrazoxane, embelin, CAN and NDGA

In addition to the aforementioned mechanisms, we further detected whether these inhibitors affected the secondary structure of NDM-1 by CD spectroscopy. In the absence of inhibitors, the NDM-1 contains 52.5% α-helix and 22.0% turn conformations. However, following treatment of NDM-1 with 32 μg/mL of the inhibitors, the percentage of α-helix conformation was reduced to 0%, 18.8%, 0% and 17.2% for dexrazoxane, embelin, CAN and NDGA, respectively (**Figure 6A**). The percentage of turn conformation was increased to 76.7%, 54.3%, 70.3% and 46.1%, respectively. Additionally, NDGA-treated NDM-1 maintained a highly consistent conformational composition with untreated NDM-1, and only the proportion of different conformations is changed, while the conformation of dexrazoxane, embelin or CAN-treated NDM-1 (including heat inactivated NDM-1) changed dramatically compared with that of untreated NDM-1 (**Figure 6A**). With the addition of inhibitors, the negative ellipticity of the CD spectrum of NDM-1 decreased, and the negative peak at 222 nm was shifted to higher wavenumbers accompanied by a shortened amplitude (**Figure 6B**). Such conformational changes may be attributed to the direct interaction between inhibitors and NDM-1 or the indirect influence of zinc ions depletion within NDM-1 caused by the inhibitors. However, the currently available structural biology data did not clearly clarify the importance of zinc ion on the structural stability of NDM-1. Thus, our results indicated that treatment with dexrazoxane, embelin, CAN and NDGA measurably altered the secondary structure of NDM-1.

**Figure 6.**
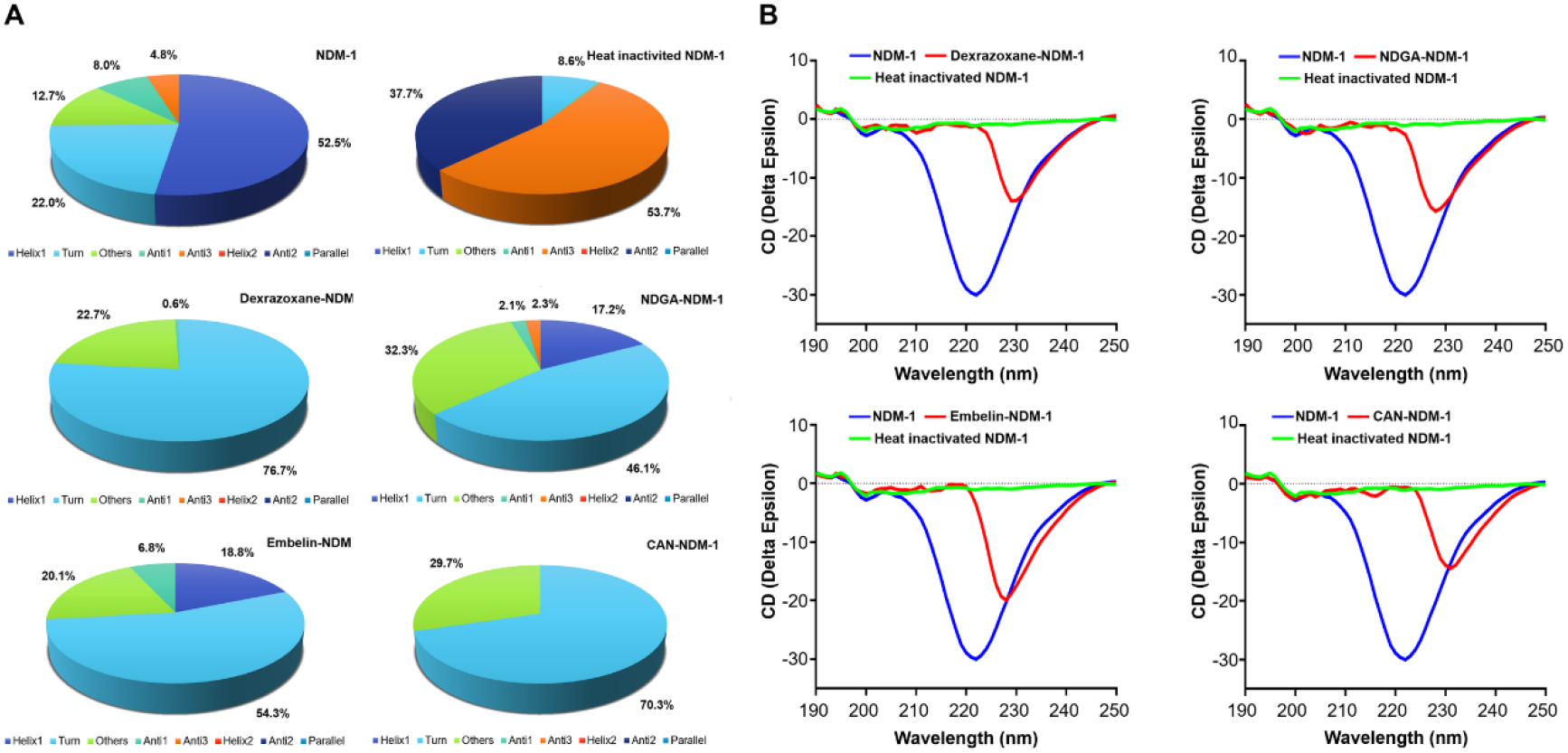
Inhibitors alter the secondary structure of NDM-1. (**A**) Secondary structure composition of NDM-1 in the presence or absence of 32 μg/mL of the inhibitors as determined by CD spectra. **(B)** Calculated CD spectrum with dexrazoxane, NDGA, embelin and CAN on NDM-1. Comparison between NDM-1 (blue), NDM-1 treated with inhibitor (red) and heat inactivated NDM-1 (green) at 70 °C for 30 min. The wavelength for CD spectroscopy was set as 190-250 nm.

### Dexrazoxane and embelin reduce the protein stability of NDM-1

In addition to the direct inhibition of NDM-1, we further assessed whether these compounds affect the production of NDM-1. Bacterial strains harbouring plasmid-borne *bla*_NDM-1_ were treated with increasing concentrations of inhibitors, and the total cell lysates were subsequently subjected to immunoblotting to determine the protein level of NDM-1. We found that the amount of NDM-1 was remarkably reduced in response to 32 μg/mL of dexrazoxane and embelin but not NDGA or CAN (**Figure 7A** and **Figure 7- figure supplement 1A**). The reduced level of NDM-1 did not result from the loss of plasmid stability by the effect of dexrazoxane and embelin (**Figure 7- figure supplement 2**). It was previously reported earlier that zinc ion depletion could accelerate the degradation of MBLs in bacteria [22]. Hence, the decreased amount of NDM-1 in the tested bacterial isolates could be the consequence of metal ion removal by dexrazoxane and embelin. Indeed, we observed a restored amount of NDM-1 in the sample with an addition of excessive zinc ions in bacteria treated with 32 μg/mL of dexrazoxane and embelin (**Figure 7B** and **Figure 7- figure supplement 1B**). CAN, another metal ion chelator, was not observed detected to reduce the amount of NDM-1 at the tested concentration, probably due to its weaker chelating capability or relatively poor permeability to bacterial cells, which failed to efficiently remove zinc ions associated with NDM-1. Taken together, our data suggested that deletion of zinc ions by dexrazoxane and embelin could induce the degradation of NDM-1.

**Figure 7.**
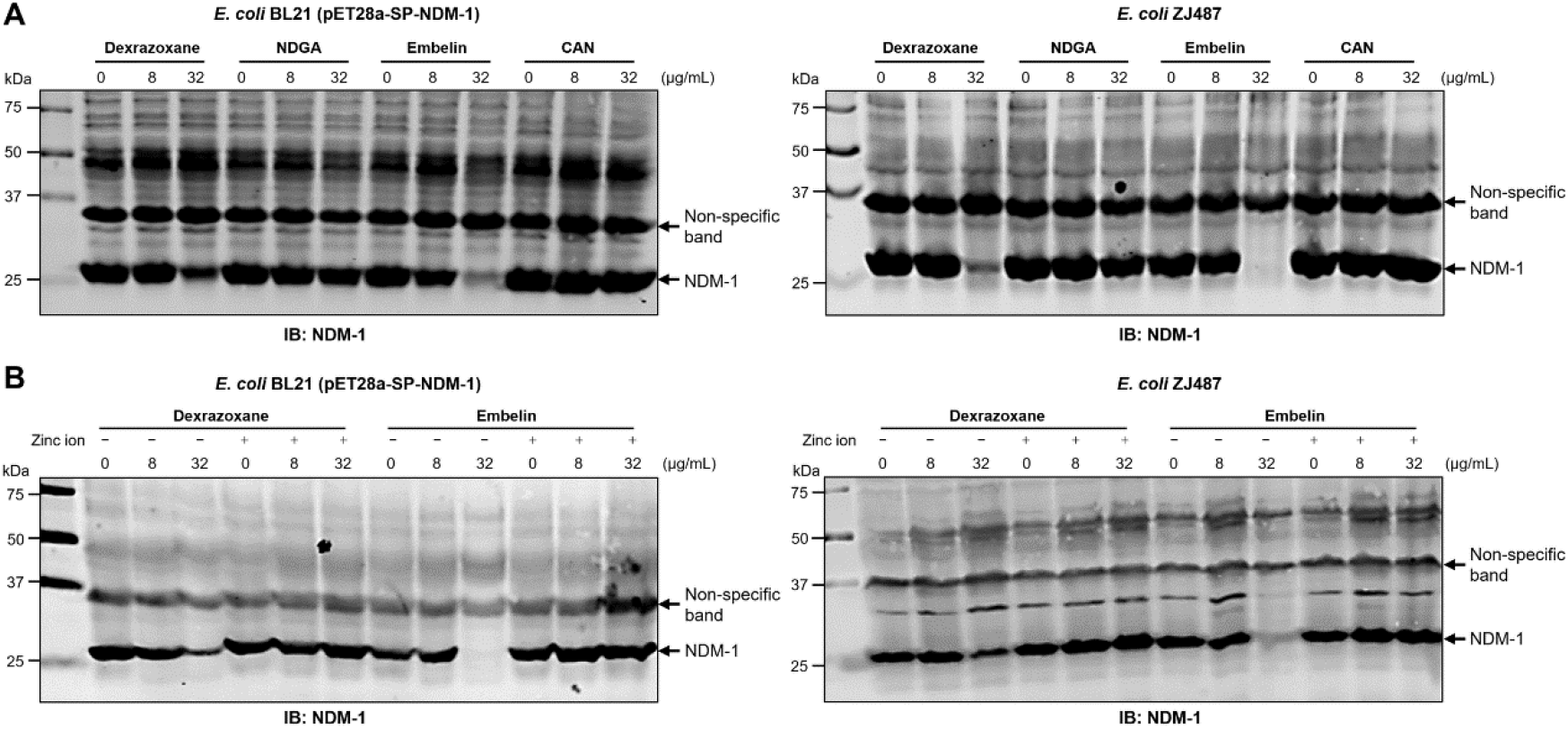
Dexrazoxane and embelin induce NDM-1 degradation via metal ion depletion manner. (**A**) NDM-1 levels in *E. coli* strains BL21 (pET28a-SP-NDM-1) and ZJ487 treated with the indicated concentrations of inhibitors. (**B**) The addition of 500 μM of zinc ions suppresses the degradation of NDM-1 resulting from dexrazoxane and embelin treatment. Total proteins of bacteria cultured in the presence or absence of inhibitors and additional zinc ions were separated by SDS-PAGE and probed with NDM-1 specific antibody. The non-specific band was used as an internal loading control. The blots shown are one representative of three independent experiments with similar observations.

### Dexrazoxane, embelin, CAN and NDGA restore meropenem activity *in vivo*

Given the potent inhibition of dexrazoxane, embelin, CAN and NDGA on the enzymatic activity of NDM-1, as well as their excellent *in vitro* synergistic bactericidal activity against NDM-1 producing bacterial pathogens when used in combination with meropenem, we further investigated the potential application of these inhibitors to overcome carbapenem resistance *in vivo* and restore the treatment efficacy of meropenem in the clinical settings. We established three mouse infection models to assess the *in vivo* therapeutic effects of the combined therapy. In the lethal systemic infection model, all mice infected with *E. coli* ZJ487 treated with PBS or meropenem monotherapy died within 36 h post-infection. However, the combined therapy of meropenem with the inhibitors resulted in 33.33%, 50%, 66.67% and 33.33% survival for dexrazoxane, NDGA, embelin and CAN, respectively (**Figure 8A**). Surprisingly, embelin alone can also increase the survival rate of mice to 33%, which may be due to its unexplored pharmacological actions on either mammalian hosts or bacteria cells.

**Figure 8.**
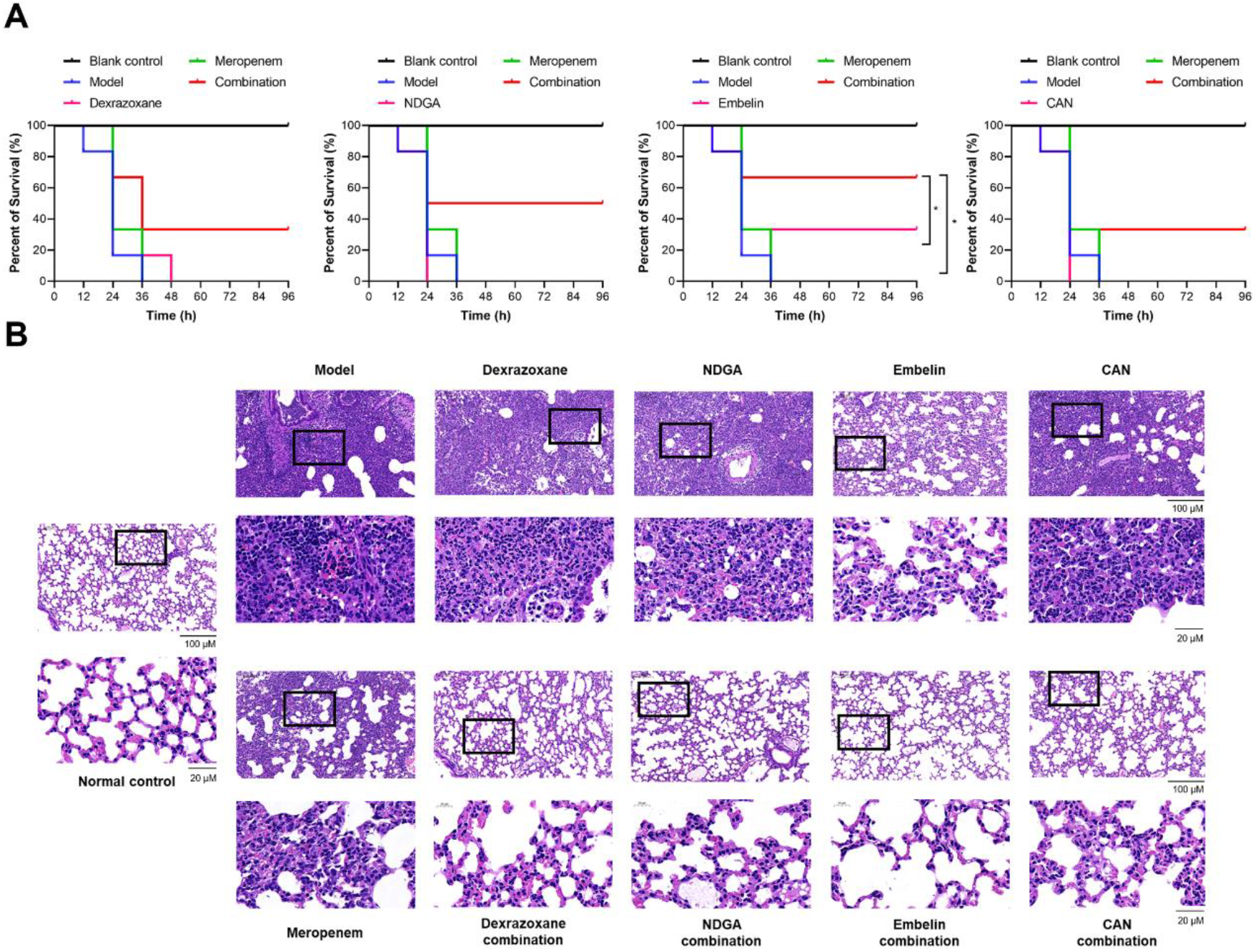
Dexrazoxane, NDGA, embelin and CAN rescue meropenem efficacy *in vivo*. Mice were challenged with *E. coli* ZJ487 and then treated with inhibitors (80 mg/kg), meropenem (10 mg/kg), inhibitors (80 mg/kg) combined with meropenem (10 mg/kg) (combination), or DMSO (model). Mice without bacterial infection were included as a blank control. (**A**) The combination of individual inhibitors and meropenem significantly increased the survival rate of mice intraperitoneally infected with *E. coli* ZJ487 compared with meropenem monotherapy. Mice were randomly allocated into different groups and each group contained 6 mice (n=6). (**B**) The combination of individual inhibitors and meropenem alleviated histopathological injury of mice in a mouse pneumonia model compared with meropenem monotherapy. Lungs of infected mice were collected 72 h post-infection and stained with haematoxylin and eosin. Histopathological changes were visualized under a digital slide scanner. Mice were randomly allocated into different groups and each group contained 6 mice (n=6). The experimental results shown in panels A and B are one representative from three independent experiments. * indicates *P* < 0.05 as determined by log-rank (Mantel-Cox) test.

The therapeutic advantages of the combined therapy were also supported by the mouse thigh muscle infection model. The bacterial load was reduced significantly in infected mice treated with meropenem and the individual NDM-1 inhibitors (**Figure 8- figure supplement 1A**). Moreover, we tested the treatment efficacy of the combination therapy in a mouse pneumonia infection model. Co-therapy of mice with meropenem and NDM-1 inhibitors resulted in strikingly lower bacterial counts in the lungs (**Figure 8- figure supplement 1B**) and as well as the marked remission of pulmonary inflammation as evidenced by less inflammatory factor infiltration in the alveolar space (**Figure 8B**). Together, these observations confirmed the potential utilization of dexrazoxane, embelin, CAN and NDGA to rescue carbapenem activity *in vivo* against infections caused by NDM-1 producing bacterial pathogens.

## Discussion

Carbapenems are still the last-resort antibiotics to control serious infectious diseases caused by MDR gram-negative bacterial pathogens. However, the development of carbapenem resistance mediated by carbapenemase has greatly limited the clinical use of carbapenems. Therefore, it is urgent to develop novel treatment strategies to address infections caused by carbapenem resistant bacteria. In recent decades, many scientific groups have concentrated on the identification of novel effective β-lactamase inhibitors, and fortunately resulting in significant progress [23]. The most striking achievements are the discovery of diazabicyclooctanones (DBOs), non-β-lactam β-lactamase inhibitors [24]. Recently, two DBOs (avibactam and relebactam) have been approved for clinical use in combination with β-lactams such as ceftazidime, imipenem and cilastatin [25–27]. Moreover, two novel DBO-type inhibitors, zidebactam and nacubactam, which have high affinity for penicillin-binding protein 2 and potent inhibition of β-lactamase, are under investigation in clinical trials combined with β-lactams [28–30]. In addition to DBOs, another type of non-β-lactam β-lactamase inhibitor, vaborbactam, which contains a cyclic boronic acid pharmacophore, was available on the market and used in conjunction with meropenem [31]. Obviously, the clinical application of DBOs and vaborbactam expanded the therapeutic options for lethal diseases by gram-negative bacteria. However, both DBOs and vaborbactam are narrow-spectrum inhibitors and are only active towards SBLs but not MBLs [24, 32]. Hence, the identification of inhibitors against MBLs is highly urgent and represents the major challenge in the field of β-lactamase inhibitor development. A great number of compounds such as thiols, thioesters, azolylthioacetamide, carboxylic acids, cyclic boronates and aspergillomarasmine A showed activity to inhibit MBLs by diverse mechanisms [19, 33]. Despite of the worldwide academic efforts, no MBLs inhibitors are close to clinical use.

Compared to the *de novo* development of an entirely new drug for a specific disease, drug repurposing, which aims to identify novel applications for existing drugs, has gained increasing interest during the last decade [34, 35]. This strategy exhibits significant advantages since the repurposed drugs already have pharmacokinetic and safety assessment data, which can greatly reduce the timeline and cost of development. Here we employed the repurposing approach and found four NDM-1 inhibitors from the FDA-approved drugs. Dexrazoxane is used mainly as an anti-tumor adjuvant drug to alleviate the cardiotoxicity induced by anthracycline [36]; NDGA is an antioxidant applied mostly in the food industry [37]; embelin, a natural compound isolated from *Embeliaribes*, is a well-known antagonist for X-linked inhibitor of apoptosis protein (XIAP) used for the treatment of various cancers [38]; CAN is a potent blocker of angiotens in II receptor approved to treat hypertension in adults [39]. These four inhibitors showed broad-spectrum inhibitory effects on MBLs either by chelating zinc ions (dexrazoxane, embelin and CAN) or by directly engaging MBLs (NDGA). When used in combination with meropenem, these inhibitors displayed potent synergistic bactericidal effects on MBLs-producing bacteria *in vitro*. In addition, combined therapy with meropenem and inhibitors also exhibits synergistic treatment advantages against infections induced by bacteria expressing NDM-1.

It was noted that an earlier study also reported embelin as an NDM-1 inhibitor from an enzymatic-based screening of naturally occurring chemicals [40]. Here, we further demonstrated that embelin functions as a potent chelating agent to directly deplete zinc ions from NDM-1, and further clarified its potential application in combined therapy with meropenem against MBLs-harbouring bacteria in various animal infection models.

## Conclusion

Our data demonstrated that the FDA-approved dexrazoxane, embelin, CAN and NDGA possess excellent synergistic activity with meropenem against carbapenem-resistant bacteria mediated by MBLs both *in vitro* and *in vivo*. Our observations together with the established safety assessments indicate the great potential of repurposing these drugs as antibiotic adjuvants to fight lethal infections caused by MBLs-producing bacterial pathogens.

## Acknowledgements

The authors thank Professor Yonghong Xiao (Zhejiang University, China) for strains *E. cloacae* 20710 and 20712, and thank Dr. Zhimin Guo (the First Hospital of Jilin University, China) for *A. baumannii* 21. This work was supported by the National Key Research and Development Program of China (2021YFD1801000), the National Natural Science Foundation of China (grant nos. 81861138046 and 32172912).

---

**Table S1.**
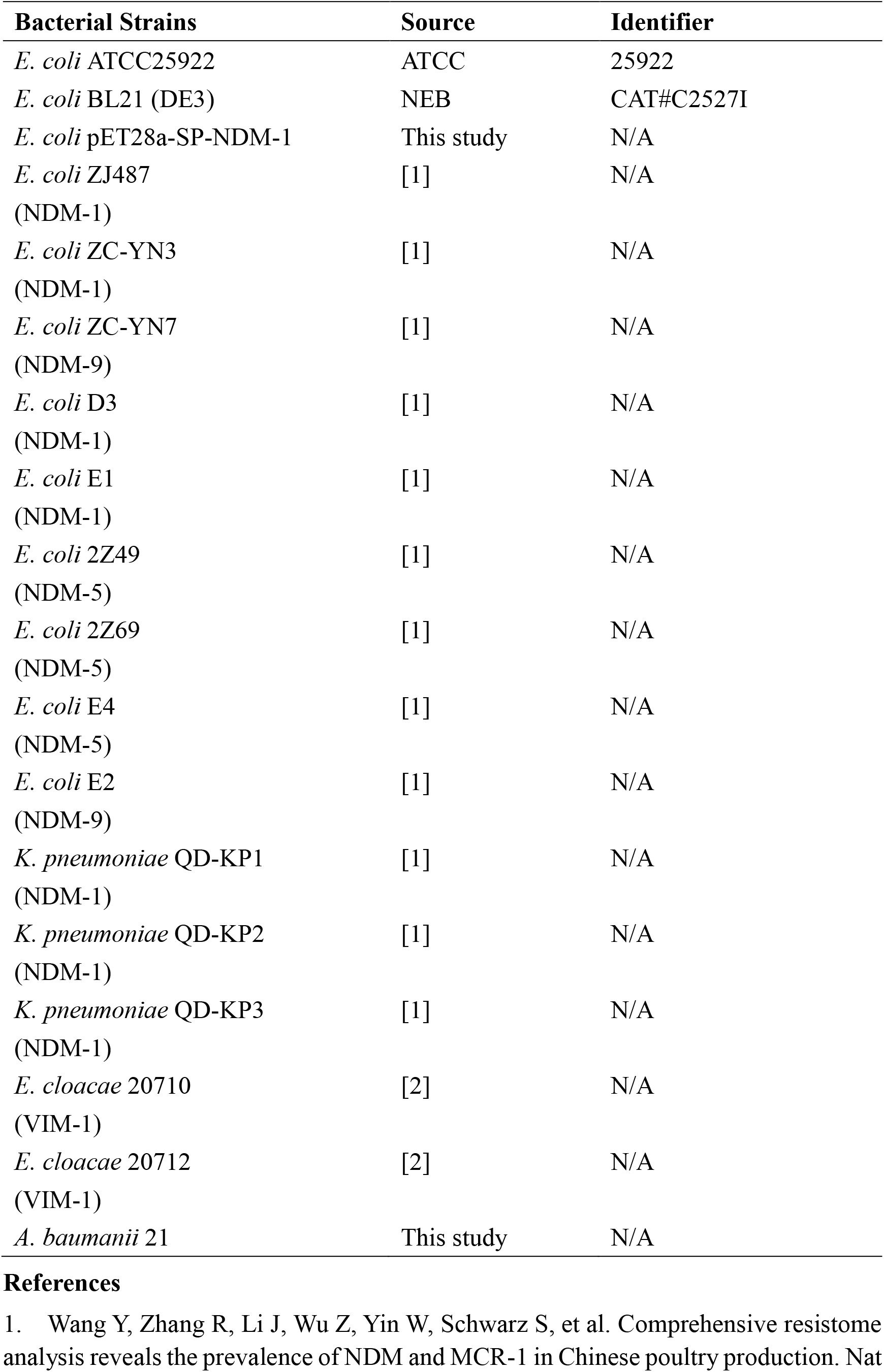
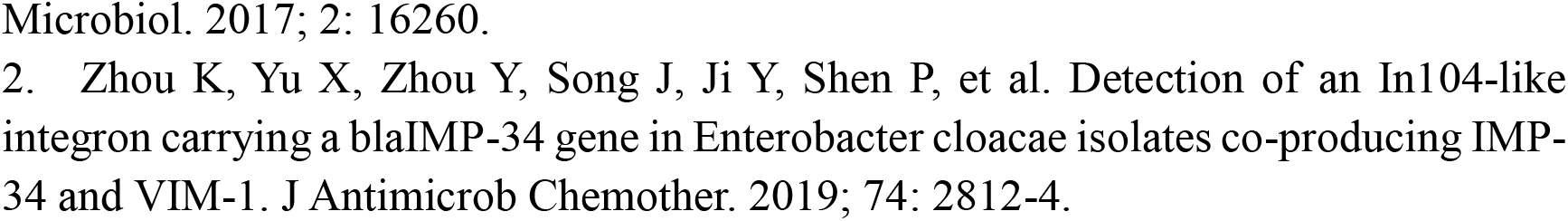
Bacterial strains used in this study.

**Table S2.** Plasmids used in this study.

| **Plasmids** | **Source** | **Identifier** |
| --- | --- | --- |
| pET28a | Novagen | CAT#69864 |
| pETSUMO | Invitrogen | CAT#K300 |
| pET28a-NDM-1 | This study | N/A |
| pETSUMO-KPC-2 | This study | N/A |
| pETSUMO-VIM-1 | This study | N/A |
| pETSUMO-IMP-1 | This study | N/A |
| pETSUMO-OXA-48 | This study | N/A |
| pET28a-NDM-3 | This study | N/A |
| pET28a-NDM-9 | This study | N/A |
| pET28a-NDM-1<sub>I35A</sub> | This study | N/A |
| pET28a-NDM-1<sub>K211A</sub> | This study | N/A |
| pET28a-NDM-1<sub>H250A</sub> | This study | N/A |

**Table S3.**
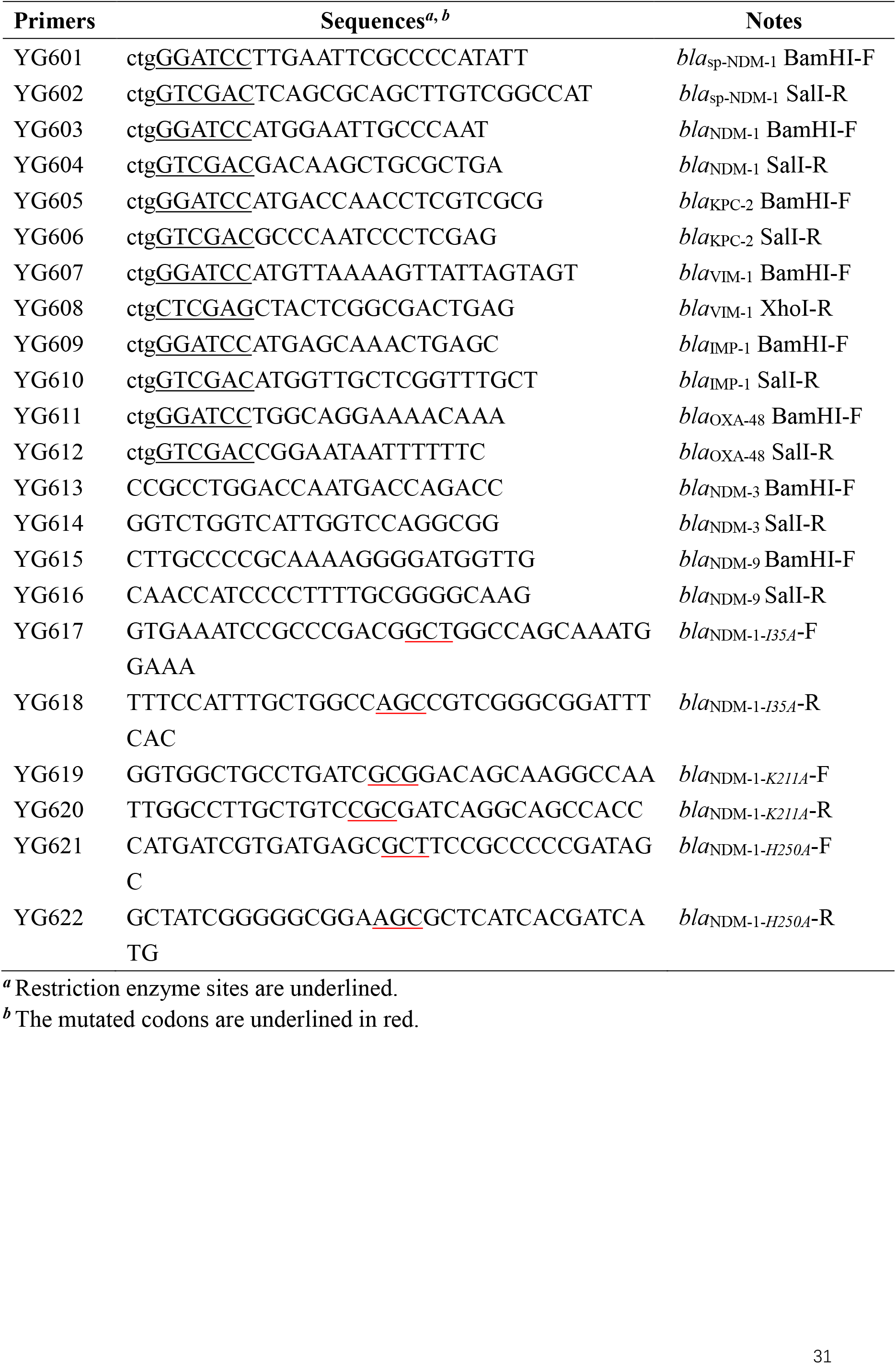
Primers used in this study.

**Table S4.**
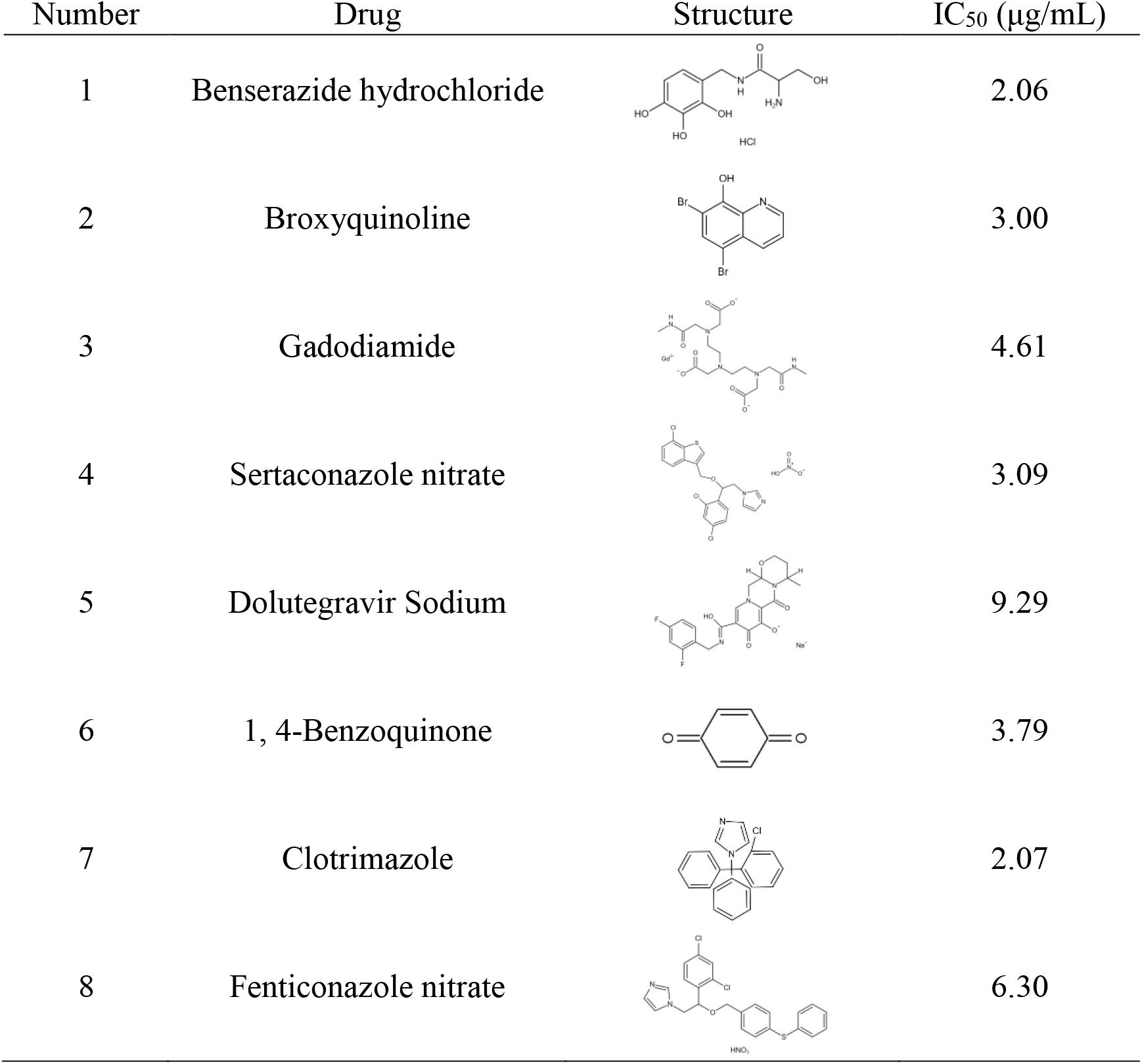
IC_50_ of the remaining active inhibitors on NDM-1.

| Number | Drug | Structure | IC<sub>50</sub> (μg/mL) |
| --- | --- | --- | --- |
| 1 | Benserazide hydrochloride |  | 2.06 |
| 2 | Broxyquinoline |  | 3.00 |
| 3 | Gadodiamide |  | 4.61 |
| 4 | Sertaconazole nitrate |  | 3.09 |
| 5 | Dolutegravir Sodium |  | 9.29 |
| 6 | 1, 4-Benzoquinone |  | 3.79 |
| 7 | Clotrimazole |  | 2.07 |
| 8 | Fenticonazole nitrate |  | 6.30 |

**Table S5.**
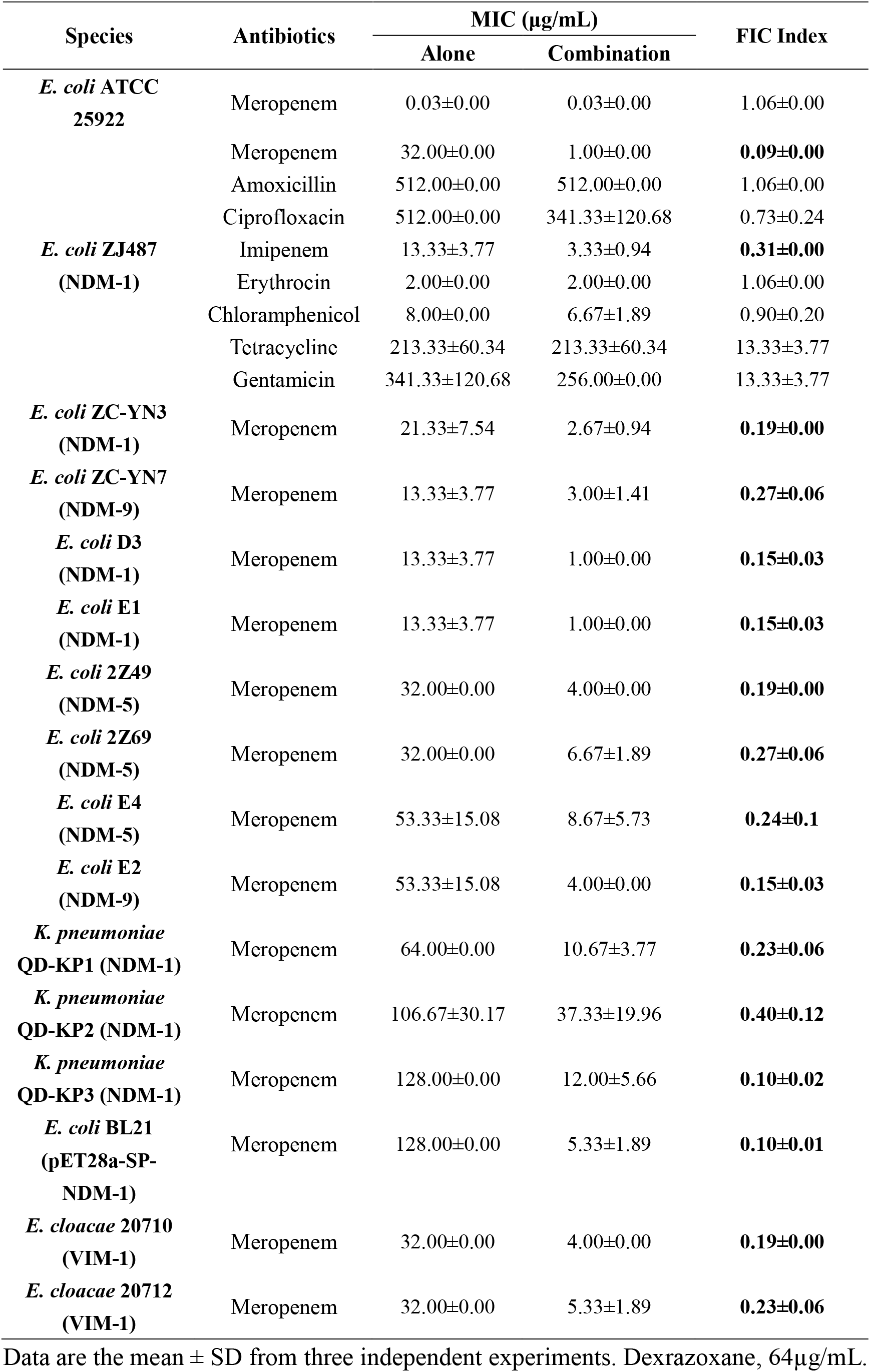
Synergistic antibacterial effect of meropenem in combination with dexrazoxane on tested strains.

**Table S6.**
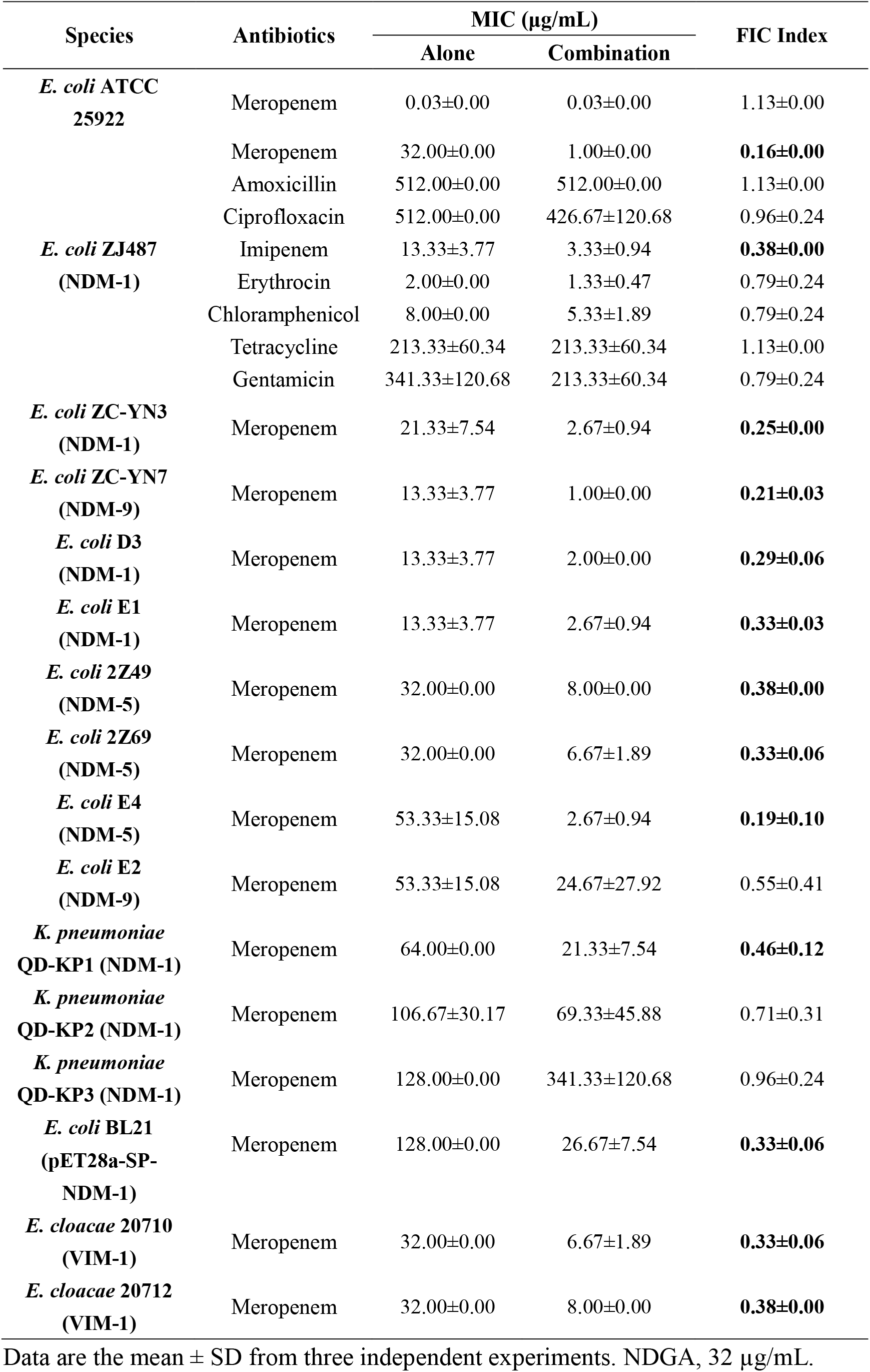
Synergistic antibacterial effect of meropenem in combination with NDGA on tested strains.

**Table S7.**
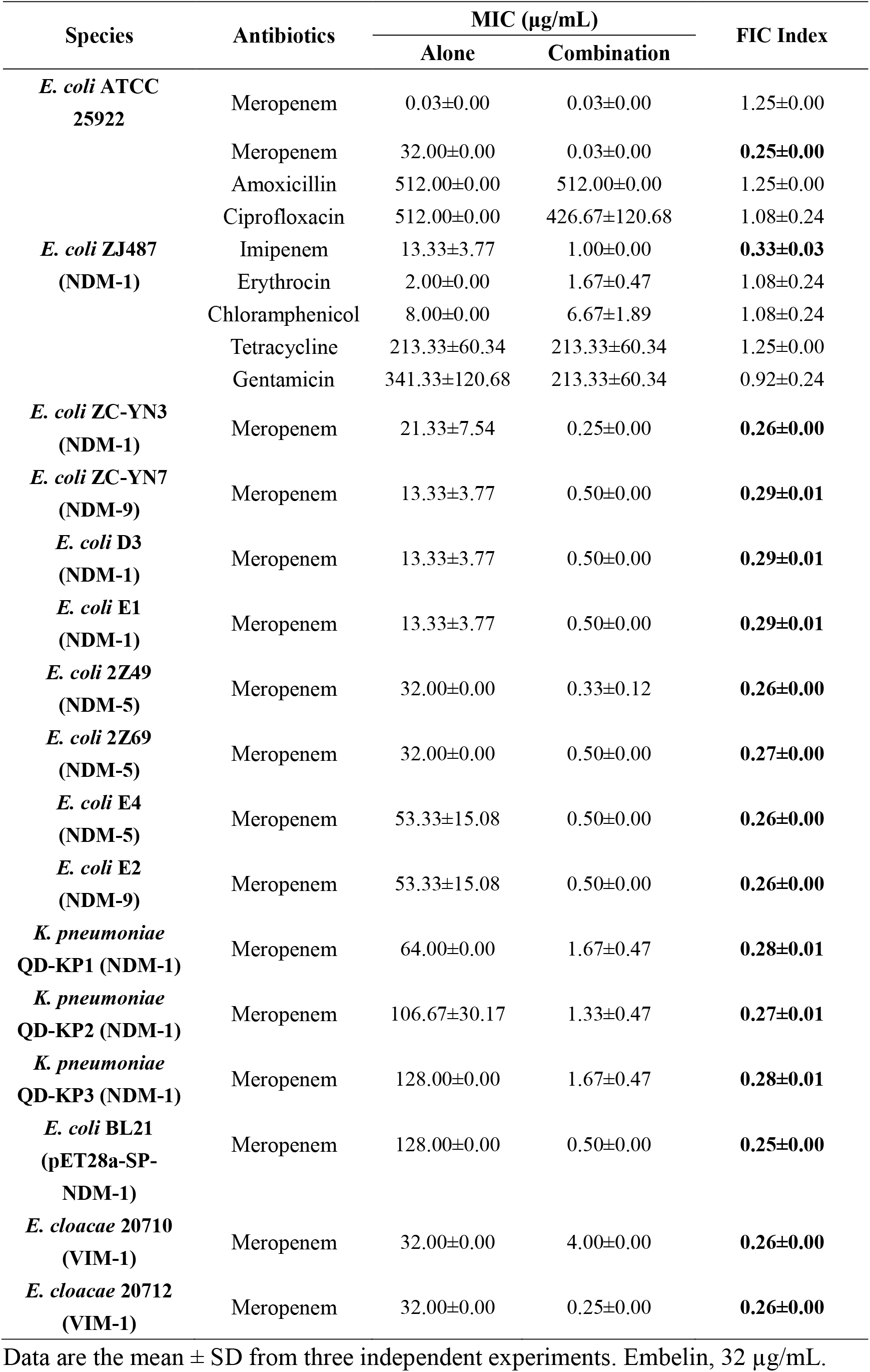
Synergistic antibacterial effect of meropenem in combination with embelin on tested strains.

**Table S8.**
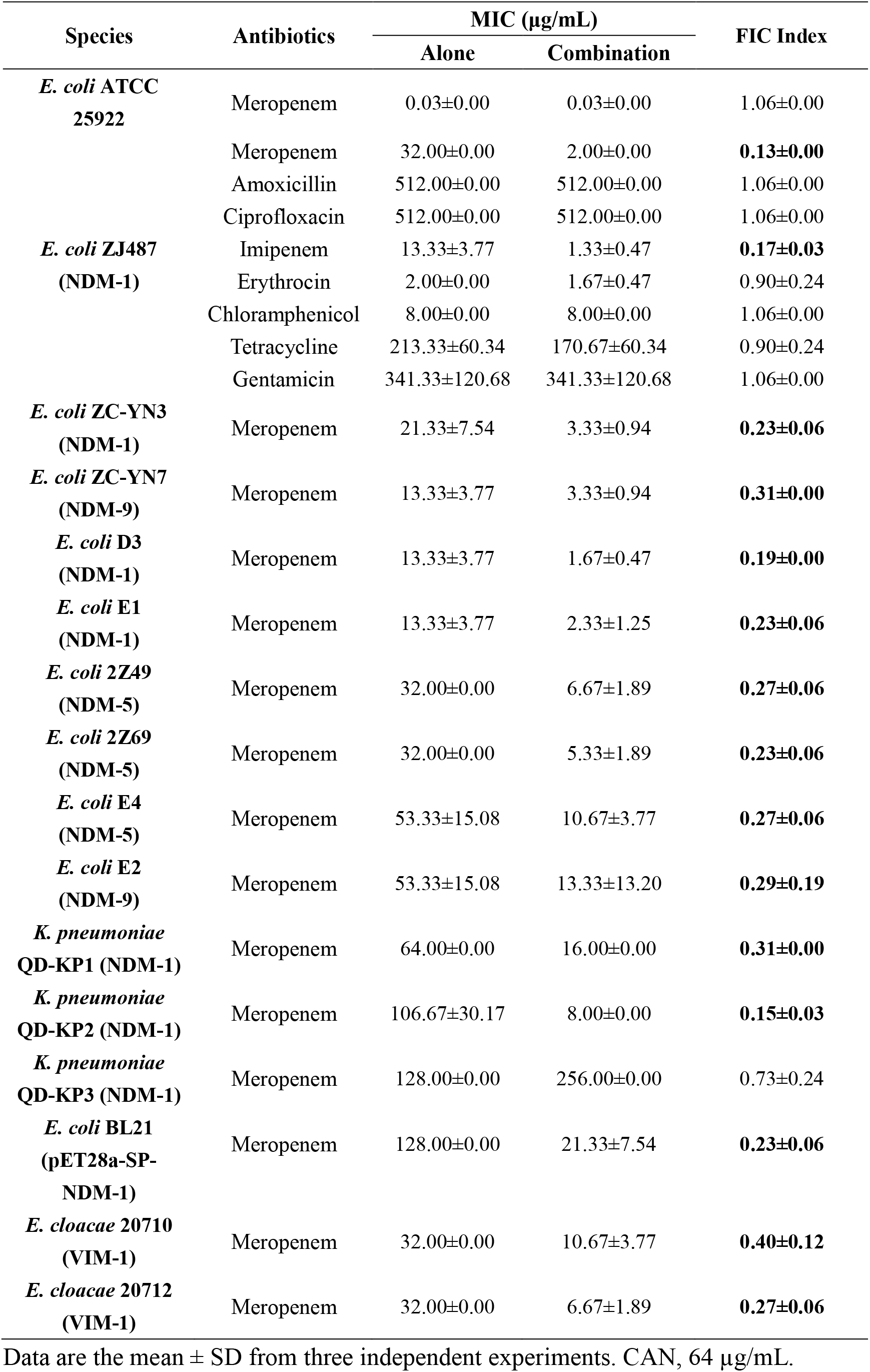
Synergistic antibacterial effect of meropenem in combination with CAN on tested strains.

**Figure 2- figure supplement 1.**
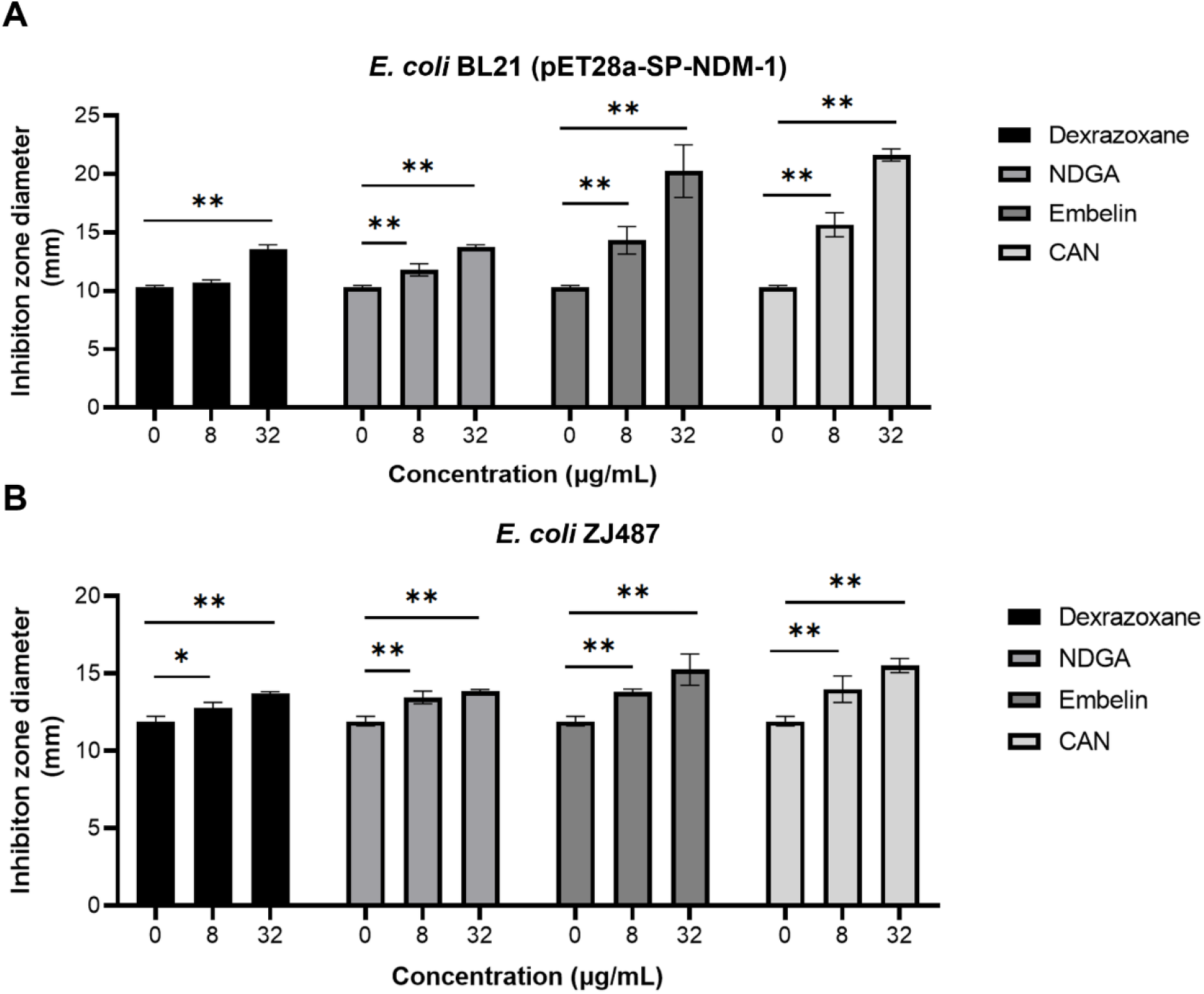
Dexrazoxane, NDGA, embelin and CAN rescued the activity of meropenem *in vitro* (related to **Figure 2B**). The inhibition zone diameter of meropenem disks supplemented with 0 µg/mL, 8 µg/mL or 32 µg/mL of dexrazoxane, NDGA, embelin and CAN against *E. coli* strains BL21 (pET28a-SP-NDM-1) **(A)** and ZJ487 **(B)**. The data shown are the mean ± SD from three independent experiments. * indicates *P* < 0.05 and ** indicates *P* < 0.01 by Student’s *t*-test.

**Figure 3- figure supplement 1.**
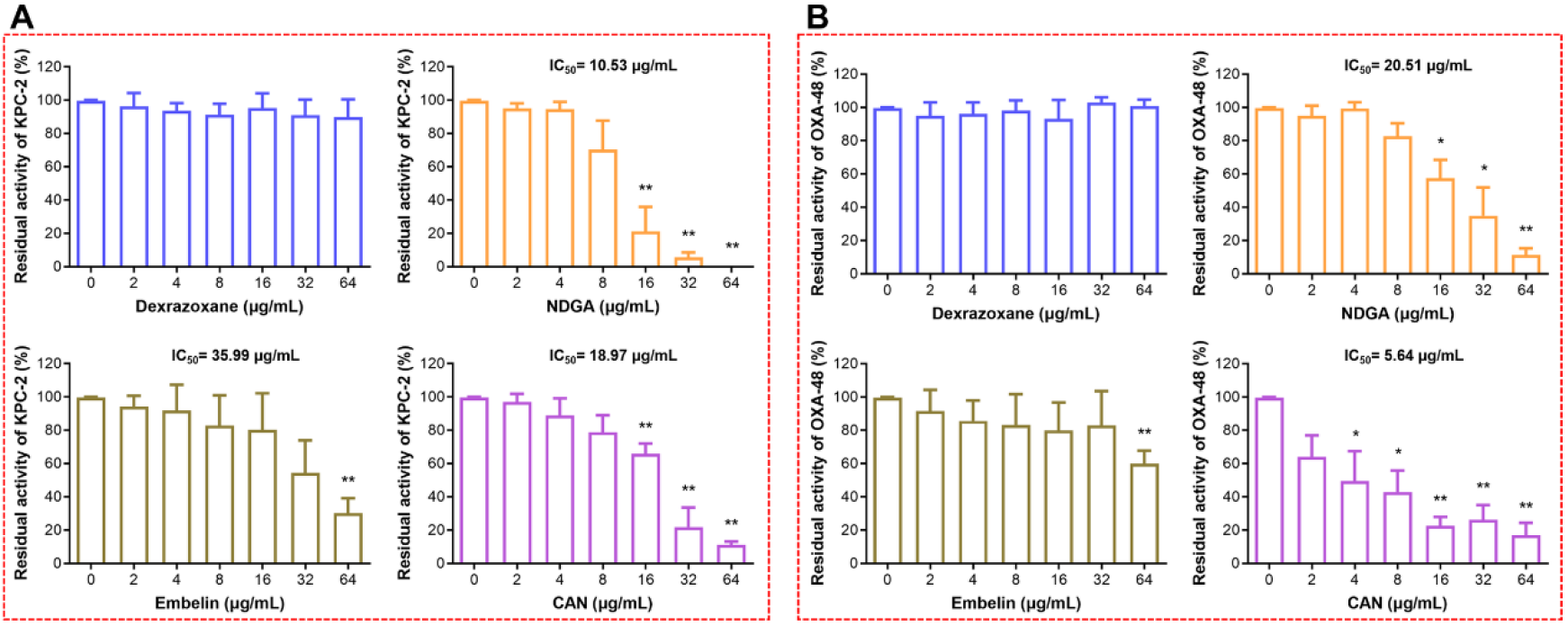
Effects of dexrazoxane, NDGA, embelin and CAN on the activity of KPC-2 and OXA-48. Inhibition of KPC-2 (**A**) and OXA-48 (**B**) by dexrazoxane, NDGA, embelin and CAN. Data represent the mean ± SD from three independent experiments. * indicates *P* < 0.05 and ** indicates *P* < 0.01 by Student’s *t*-test.

**Figure 7- figure supplement 1.**
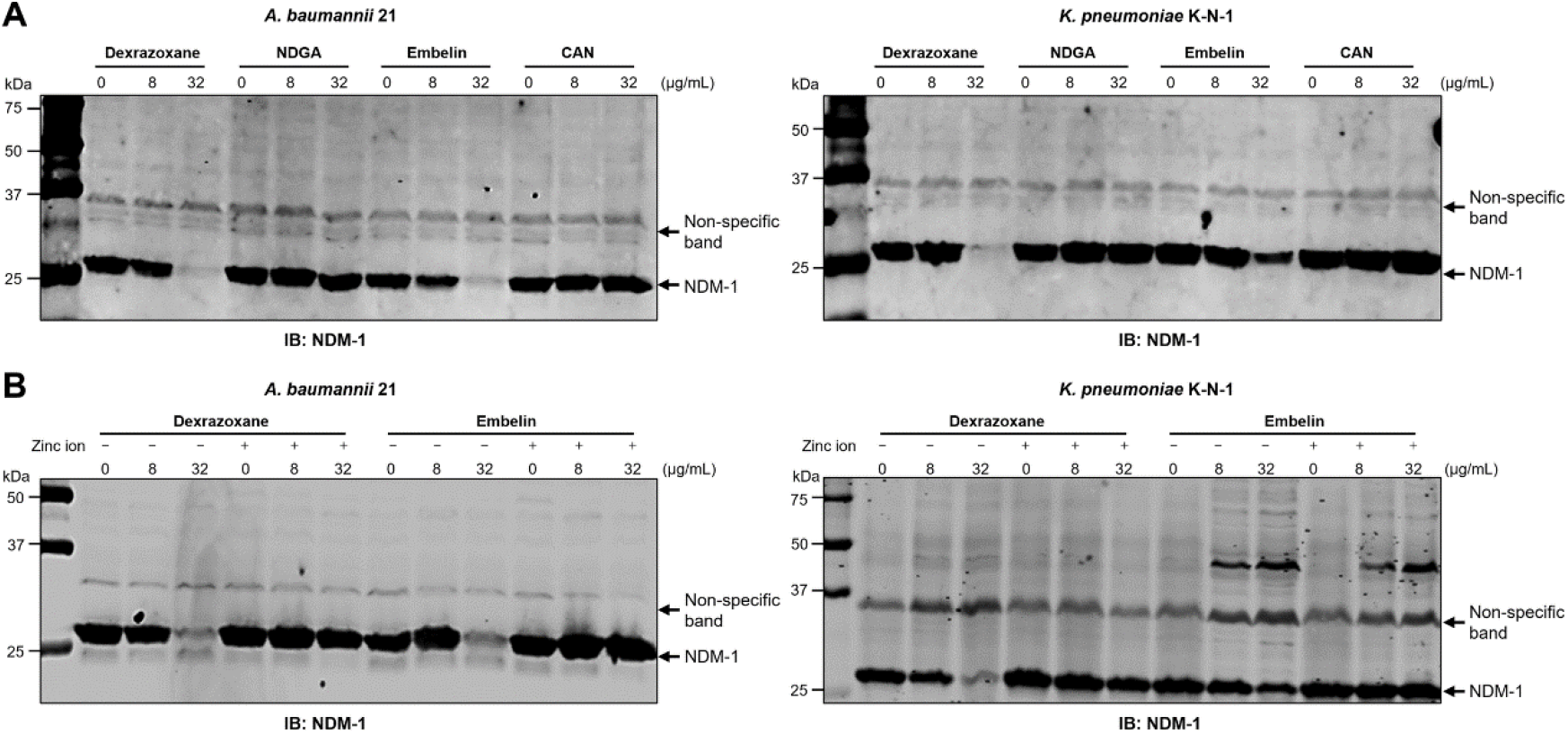
Dexrazoxane and embelin induce NDM-1 degradation via metal ion depletion manner. (**A**) NDM-1 levels in *A. baumannii 21* and *K. pneumoniae* K-N-1 treated with the indicated concentrations of inhibitors. (**B**) The addition of 500 μM of zinc ions suppresses the degradation of NDM-1 resulting from dexrazoxane and embelin treatment. Total proteins of bacteria cultured in the presence or absence of inhibitors and additional zinc ions were separated by SDS-PAGE and probed with NDM-1 specific antibody. The non-specific band was used as an internal loading control. The blots shown are one representative of three independent experiments with similar observations.

**Figure 7- figure supplement 2.**
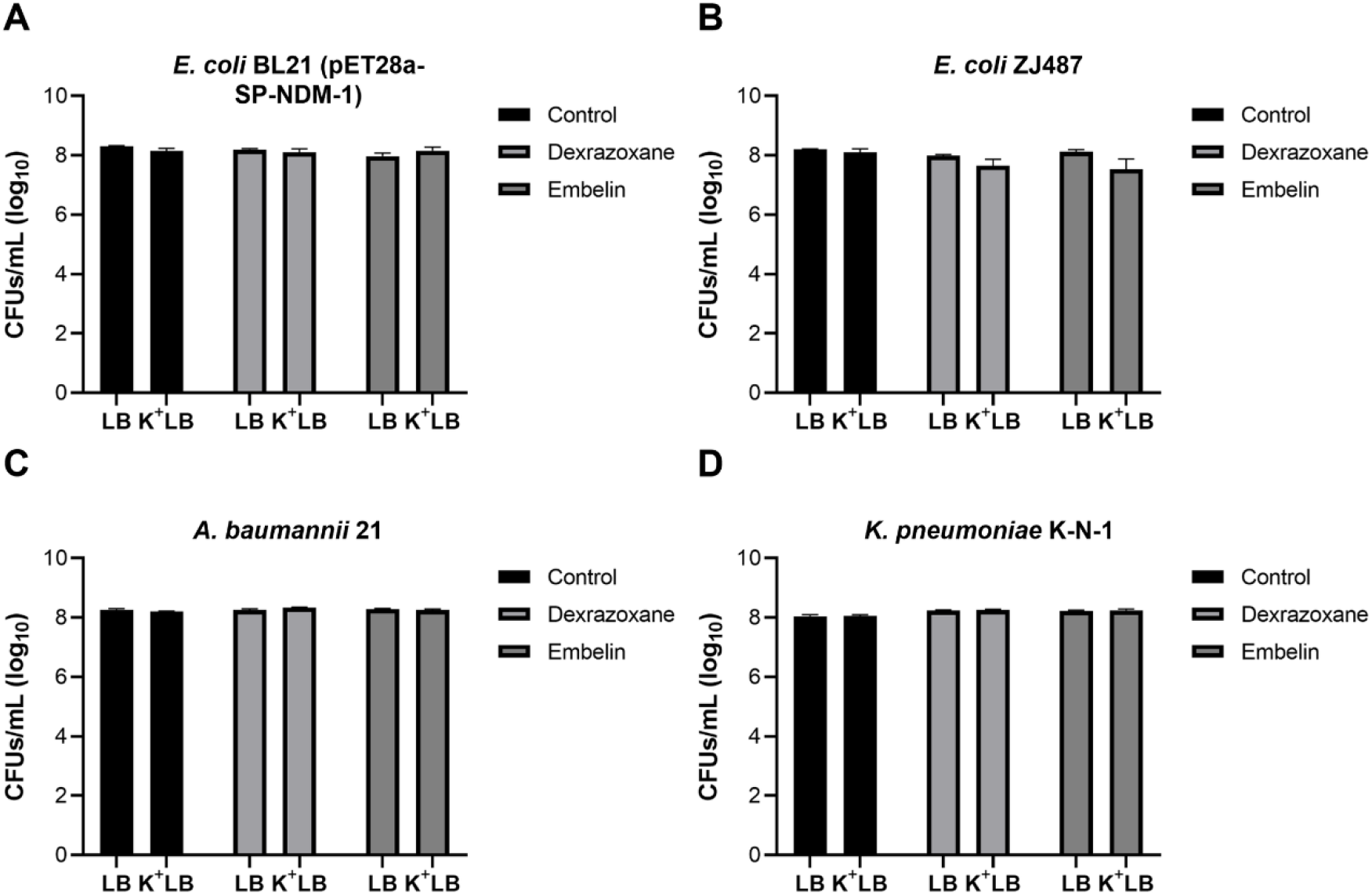
Dexrazoxane and embelin do not affect plasmid stability. Plasmid stability of *E. coli* BL21 (pET28a-SP-NDM-1) (**A**), *E. coli* ZJ487 (**B**), *A. baumannii* 21 (**C**) or *K. pneumoniae* K-N-1 (**D**) following culture with 32 μg/mL of dexrazoxane or embelin. The data shown are the mean ± SD from three independent experiments.

**Figure 8- figure supplement 1.**
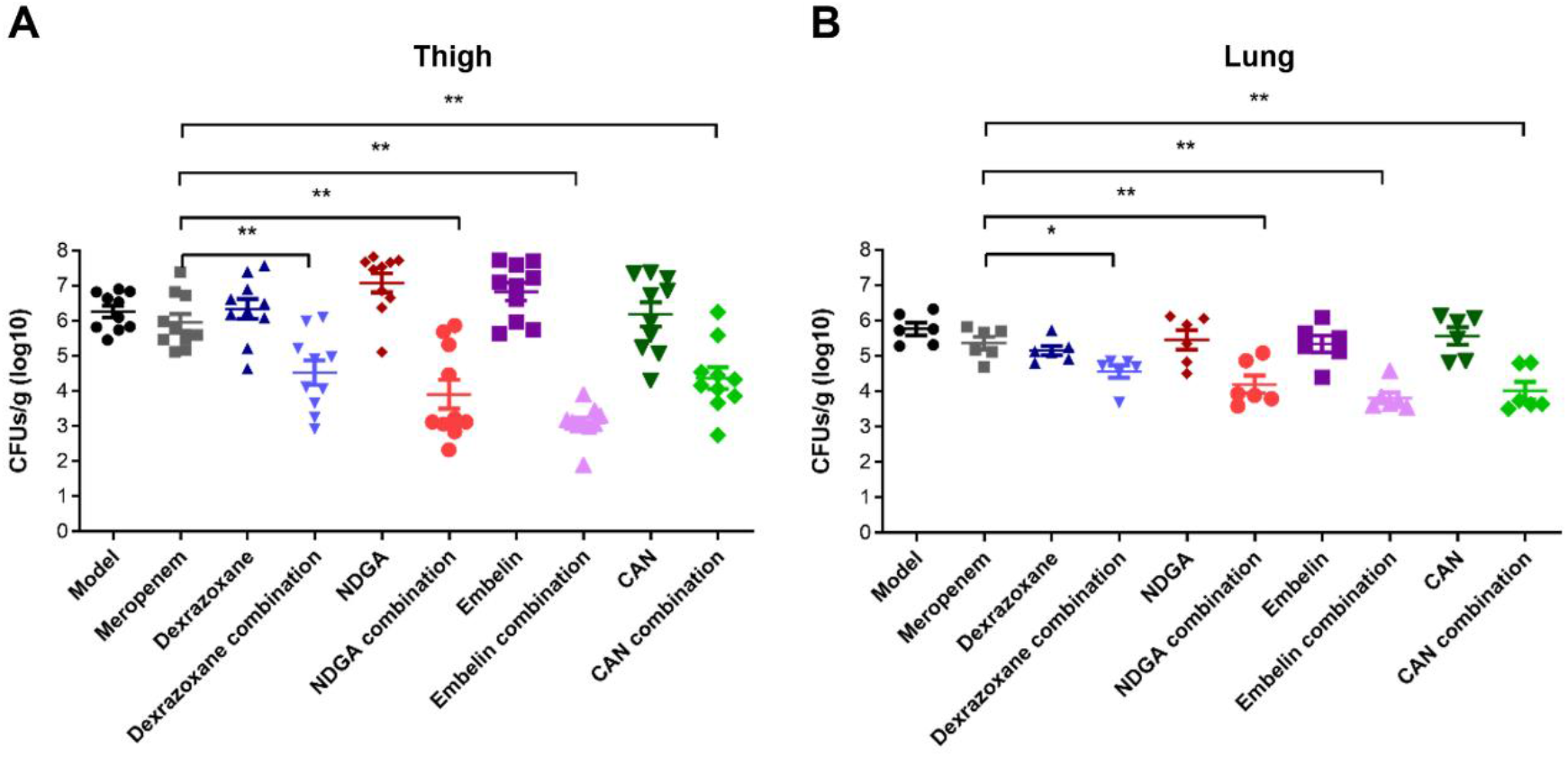
Dexrazoxane, NDGA, embelin and CAN rescue meropenem activity *in vivo*. (**A**) The combination of meropenem with dexrazoxane, NDGA, embelin or CAN decreased the bacterial load in the mouse thigh infection model compared with the meropenem monotherapy. Mice were randomly allocated into different groups and each group contains 10 mice (n=10). (**B**) The combination of meropenem with dexrazoxane, NDGA, embelin or CAN reduced the bacterial load of the lungs in the mouse pneumonia model compared with the meropenem monotherapy (related to Figure 8B). Mice were randomly allocated into different groups and each group contains 6 mice (n=6). The data shown are one representative of three independent experiments. * indicates *P* < 0.05 and ** indicates *P* < 0.01 by Mann–Whitney U test.

## References

1. De Oliveira DMP, Forde BM, Kidd TJ, Harris PNA, Schembri MA, Beatson SA, et al. Antimicrobial Resistance in ESKAPE Pathogens. Clin Microbiol Rev. 2020; 33.

2. de Kraker ME, Stewardson AJ, Harbarth S. Will 10 Million People Die a Year due to Antimicrobial Resistance by 2050? PLoS Med. 2016; 13: e1002184.

3. McKenna M. Antibiotic resistance: the last resort. Nature. 2013; 499: 394–6.

4. Zequinao T, Telles JP, Gasparetto J, Tuon FF. Carbapenem stewardship with ertapenem and antimicrobial resistance-a scoping review. Rev Soc Bras Med Trop. 2020; 53: e20200413.

5. Patel G, Bonomo RA. “Stormy waters ahead”: global emergence of carbapenemases. Front Microbiol. 2013; 4: 48.

6. Nordmann P, Poirel L. Epidemiology and Diagnostics of Carbapenem Resistance in Gram-negative Bacteria. Clin Infect Dis. 2019; 69: S521–S8.

7. Queenan AM, Bush K. Carbapenemases: the versatile beta-lactamases. Clin Microbiol Rev. 2007; 20: 440–58, table of contents.

8. Yong D, Toleman MA, Giske CG, Cho HS, Sundman K, Lee K, et al. Characterization of a new metallo-beta-lactamase gene, bla(NDM-1), and a novel erythromycin esterase gene carried on a unique genetic structure in Klebsiella pneumoniae sequence type 14 from India. Antimicrob Agents Chemother. 2009; 53: 5046–54.

9. van Duin D, Doi Y. The global epidemiology of carbapenemase-producing Enterobacteriaceae. Virulence. 2017; 8: 460–9.

10. Castanheira M, Deshpande LM, Mendes RE, Canton R, Sader HS, Jones RN. Variations in the Occurrence of Resistance Phenotypes and Carbapenemase Genes Among Enterobacteriaceae Isolates in 20 Years of the SENTRY Antimicrobial Surveillance Program. Open Forum Infect Dis. 2019; 6: S23–S33.

11. WHO. Global Priority List of Antibiotic-Resistant Bacteria to Guide Research, Discovery, and Development of New Antibiotics. . 2020.

12. Bush K, Bradford PA. Interplay between beta-lactamases and new beta-lactamase inhibitors. Nat Rev Microbiol. 2019; 17: 295–306.

13. Pasteran F, Gonzalez LJ, Albornoz E, Bahr G, Vila AJ, Corso A. Triton Hodge Test: Improved Protocol for Modified Hodge Test for Enhanced Detection of NDM and Other Carbapenemase Producers. J Clin Microbiol. 2016; 54: 640–9.

14. Kumar N, Singh VA, Beniwal V, Pottathil S. Modified Carba NP Test: Simple and rapid method to differentiate KPC- and MBL-producing Klebsiella species. J Clin Lab Anal. 2018; 32: e22448.

15. Howard JC, Creighton J, Ikram R, Werno AM. Comparison of the performance of three variations of the Carbapenem Inactivation Method (CIM, modified CIM [mCIM] and in-house method (iCIM)) for the detection of carbapenemase-producing Enterobacterales and non-fermenters. J Glob Antimicrob Resist. 2020; 21: 78–82.

16. Zhanel GG, Lawrence CK, Adam H, Schweizer F, Zelenitsky S, Zhanel M, et al. Imipenem-Relebactam and Meropenem-Vaborbactam: Two Novel Carbapenem-beta-Lactamase Inhibitor Combinations. Drugs. 2018; 78: 65–98.

17. Liu S, Zhang J, Zhou Y, Hu N, Li J, Wang Y, et al. Pterostilbene restores carbapenem susceptibility in New Delhi metallo-beta-lactamase-producing isolates by inhibiting the activity of New Delhi metallo-beta-lactamases. Br J Pharmacol. 2019; 176: 4548–57.

18. Zhou K, Yu X, Zhou Y, Song J, Ji Y, Shen P, et al. Detection of an In104-like integron carrying a blaIMP-34 gene in Enterobacter cloacae isolates co-producing IMP-34 and VIM-1. J Antimicrob Chemother. 2019; 74: 2812–4.

19. King AM, Reid-Yu SA, Wang W, King DT, De Pascale G, Strynadka NC, et al. Aspergillomarasmine A overcomes metallo-beta-lactamase antibiotic resistance. Nature. 2014; 510: 503–6.

20. Hess B, Kutzner C, van der Spoel D, Lindahl E. GROMACS 4: Algorithms for Highly Efficient, Load-Balanced, and Scalable Molecular Simulation. Journal of chemical theory and computation. 2008; 4: 435–47.

21. Zhou Y, Guo Y, Wen Z, Ci X, Xia L, Wang Y, et al. Isoalantolactone Enhances the Antimicrobial Activity of Penicillin G against Staphylococcus aureus by Inactivating beta-lactamase during Protein Translation. Pathogens. 2020; 9.

22. Sychantha D, Rotondo CM, Tehrani K, Martin NI, Wright GD. Aspergillomarasmine A inhibits metallo-beta-lactamases by selectively sequestering Zn(2). J Biol Chem. 2021; 297: 100918.

23. Tooke CL, Hinchliffe P, Bragginton EC, Colenso CK, Hirvonen VHA, Takebayashi Y, et al. beta-Lactamases and beta-Lactamase Inhibitors in the 21st Century. J Mol Biol. 2019; 431: 3472–500.

24. de Sousa Coelho F, Mainardi JL. The multiple benefits of second-generation beta-lactamase inhibitors in treatment of multidrug-resistant bacteria. Infect Dis Now. 2021; 51: 510–7.

25. Giri P, Patel H, Srinivas NR. Review of Clinical Pharmacokinetics of Avibactam, A Newly Approved non-beta lactam beta-lactamase Inhibitor Drug, In Combination Use With Ceftazidime. Drug Res (Stuttg). 2019; 69: 245–55.

26. Campanella TA, Gallagher JC. A Clinical Review and Critical Evaluation of Imipenem-Relebactam: Evidence to Date. Infect Drug Resist. 2020; 13: 4297–308.

27. McCarthy MW. Clinical Pharmacokinetics and Pharmacodynamics of Imipenem-Cilastatin/Relebactam Combination Therapy. Clin Pharmacokinet. 2020; 59: 567–73.

28. Moya B, Barcelo IM, Bhagwat S, Patel M, Bou G, Papp-Wallace KM, et al. WCK 5107 (Zidebactam) and WCK 5153 Are Novel Inhibitors of PBP2 Showing Potent “beta-Lactam Enhancer” Activity against Pseudomonas aeruginosa, Including Multidrug-Resistant Metallo-beta-Lactamase-Producing High-Risk Clones. Antimicrob Agents Chemother. 2017; 61.

29. Livermore DM, Mushtaq S, Warner M, Vickers A, Woodford N. In vitro activity of cefepime/zidebactam (WCK 5222) against Gram-negative bacteria. J Antimicrob Chemother. 2017; 72: 1373–85.

30. Mallalieu NL, Winter E, Fettner S, Patel K, Zwanziger E, Attley G, et al. Safety and Pharmacokinetic Characterization of Nacubactam, a Novel beta-Lactamase Inhibitor, Alone and in Combination with Meropenem, in Healthy Volunteers. Antimicrob Agents Chemother. 2020; 64.

31. Wu G, Cheon E. Meropenem-vaborbactam for the treatment of complicated urinary tract infections including acute pyelonephritis. Expert Opin Pharmacother. 2018; 19: 1495–502.

32. Lomovskaya O, Sun D, Rubio-Aparicio D, Nelson K, Tsivkovski R, Griffith DC, et al. Vaborbactam: Spectrum of Beta-Lactamase Inhibition and Impact of Resistance Mechanisms on Activity in Enterobacteriaceae. Antimicrob Agents Chemother. 2017; 61.

33. Nagulapalli Venkata KC, Ellebrecht M, Tripathi SK. Efforts towards the inhibitor design for New Delhi metallo-beta-lactamase (NDM-1). Eur J Med Chem. 2021; 225: 113747.

34. Pushpakom S, Iorio F, Eyers PA, Escott KJ, Hopper S, Wells A, et al. Drug repurposing: progress, challenges and recommendations. Nat Rev Drug Discov. 2019; 18: 41–58.

35. Peyclit L, Baron SA, Rolain JM. Drug Repurposing to Fight Colistin and Carbapenem-Resistant Bacteria. Front Cell Infect Microbiol. 2019; 9: 193.

36. Popelova O, Sterba M, Haskova P, Simunek T, Hroch M, Guncova I, et al. Dexrazoxane-afforded protection against chronic anthracycline cardiotoxicity in vivo: effective rescue of cardiomyocytes from apoptotic cell death. Br J Cancer. 2009; 101: 792–802.

37. Peralta I, Marrassini C, Filip R, Alonso MR, Anesini C. Food preservation by Larrea divaricata extract: participation of polyphenols. Food Sci Nutr. 2018; 6: 1269–75.

38. Sheng Z, Ge S, Gao M, Jian R, Chen X, Xu X, et al. Synthesis and Biological Activity of Embelin and its Derivatives: An Overview. Mini Rev Med Chem. 2020; 20: 396–407.

39. Sawhney N, Patel MK, Schachter M, Hughes AD. Inhibition of proliferation by heparin and expression of p53 in cultured human vascular smooth muscle cells. J Hum Hypertens. 1997; 11: 611–4.

40. Ning NZ, Liu X, Chen F, Zhou P, Hu L, Huang J, et al. Embelin Restores Carbapenem Efficacy against NDM-1-Positive Pathogens. Front Microbiol. 2018; 9: 71.

## References

1. Wang Y, Zhang R, Li J, Wu Z, Yin W, Schwarz S, et al. Comprehensive resistome analysis reveals the prevalence of NDM and MCR-1 in Chinese poultry production. Nat Microbiol. 2017; 2: 16260.

2. Zhou K, Yu X, Zhou Y, Song J, Ji Y, Shen P, et al. Detection of an In104-like integron carrying a blaIMP-34 gene in Enterobacter cloacae isolates co-producing IMP-34 and VIM-1. J Antimicrob Chemother. 2019; 74: 2812–4.

